# Targeted bisulfite sequencing: A novel tool for the assessment of DNA methylation with high sensitivity and increased coverage

**DOI:** 10.1101/2020.05.05.078386

**Authors:** D.A. Moser, S. Müller, E.M. Hummel, A.S. Limberg, L. Dieckmann, L. Frach, J. Pakusch, V Flasbeck, M. Brüne, J. Beygo, L. Klein-Hitpass, R. Kumsta

## Abstract

DNA methylation analysis is increasingly used in stress research. Available methods are expensive, laborious and often limited by either the analysis of short CpG stretches or low assay sensitivity. Here, we present a cost-efficient next generation sequencing-based strategy for the simultaneous investigation of multiple candidate genes in large cohorts. To illustrate the method, we present analysis of four candidate genes commonly assessed in psychoneuroendocrine research: *Glucocorticoid receptor* (*NR3C1), Serotonin transporter (SLC6A4), FKBP Prolyl isomerase 5 (FKBP5)*, and the *Oxytocin receptor* (*OXTR).*

DNA methylation standards and DNA of a female and male donor were bisulfite treated in three independent trials and were used to generate sequencing libraries for 42 CpGs from the *NR3C1 1F* promoter region, 83 CpGs of the *SLC6A4* 5’ regulatory region, 5 CpGs located in *FKBP5* intron 7, and additional 12 CpGs located in a potential enhancer element in intron 3 of the *OXTR*. In addition, DNA of 45 patients with borderline personality disorder (BPD) and 45 healthy controls was assayed. Multiplex libraries of all samples were sequenced on a MiSeq system and analyzed for mean methylation values of all CpG sites using amplikyzer2 software.

Results indicated excellent accuracy of the assays when investigating replicates generated from the same bisulfite converted DNA, and very high linearity (R^2^> 0.9) of the assays shown by the analysis of differentially methylated DNA standards. Comparing DNA methylation between BPD and healthy controls revealed no biologically relevant differences.

The technical approach as described here facilitates targeted DNA methylation analysis and represents a highly sensitive, cost-efficient and high throughput tool to close the gap between coverage and precision in epigenetic research of stress-associated phenotypes.

## Introduction

Epigenetic processes have been traditionally studied for their essential role in normal cellular development, differentiation, and regulation of gene activity or function. More recently, it has been shown that the epigenome can potentially be influenced by external factors – for instance, epigenomic changes have been correlated with the exposure to various nutritional, chemical and physical risk factors (Feil and Fraga, 2012), and perturbations in epigenetic processes have been linked to different pathologies (Godfrey et al., 2015). The most widely investigated epigenetic mechanism is DNA methylation, which can occur at around 28 million CpG sites that are distributed throughout the human genome (Edwards et al., 2010).

Epigenetics became particularly relevant for psychoneuroendocrinology, and developmental psychobiology in general, with the pioneering work by the Michael Meaney group. In a rodent model, they showed for the first time that variation in the *psychosocial* environment, i.e. natural variation in maternal care, could epigenetically program the offspring’s stress response (Weaver et al., 2004). This finding was the inception of epigenetic research in the mental health field, and since then many genes have been investigated for differential DNA methylation in relation to early life stress and/or in association with stress regulation in human and in animal models. The most widely studied candidate gene is the glucocorticoid receptor gene (*NR3C1*), with most researchers focusing on the promoter of alternative exon 1F, the human orthologue of the rat’s exon 17. Associations between early life adversity and altered *NR3C1* DNA methylation in non-neuronal cells have been reported, with the majority of studies showing increased *NR3C1* promoter DNA methylation following prenatal or postnatal stress exposure (Argentieri et al., 2017); reviewed by (Turecki and Meaney, 2016). Several other candidate genes, including the *serotonin transporter gene* (*SLC6A4*; Dammann et al., 2011; Philibert et al., 2007; van IJzendoorn et al., 2010), the *FK506 binding protein 5* gene – an important regulator of glucocorticoid sensitivity (*FKBP5*; Hohne et al., 2015; Klengel et al., 2013; Tyrka et al., 2015; Zannas et al., 2016), and the *oxytocin receptor gene* (*OXTR*; Cecil et al., 2014; Gouin et al., 2017; Unternaehrer et al., 2012) have been investigated for differential DNA methylation, and associations with different stress-related phenotypes have been described.

For the quantification of DNA methylation several methods are available, varying in costs, coverage (genome-wide vs candidate gene approaches), and resolution (see Appendix 1 for overview of methods). The most comprehensive method is whole-genome bisulfite sequencing (WGBS) that offers true genome-wide coverage by sequencing the whole genome including all ∼28 million CpGs after bisulfite treatment of DNA (Cokus et al., 2008; Laurent et al., 2010; Lister et al., 2009; Urich et al., 2015). Another method – reduced representation bisulfite sequencing (RRBS; Meissner et al., 2005) – reduces costs and complexity of the analysis by performing an initial digest of DNA, which cleaves away rather uninformative DNA. This leads to a massive reduction in amount of DNA to be assayed and thus to reduced costs. Both of the aforementioned methods have in common that they are rather useful in forensic research but are still prohibitive in larger samples because of the associated costs and bioinformatics demands. The Infinium Methylation Epic Beadchip represents an affordable platform for assessing DNA methylation for up to 850.000 single CpG sites across the genome (Bibikova et al., 2011; Pidsley et al., 2016). This array-based method is most suitable as a tool to identify associations between single (or clusters of) CpG sites and phenotypes of interest, which can be directly mapped to genes. Although the Epic Beadchip only assays ∼ 3 % of all genomic CpGs, it represents a highly informative and cost efficient tool to screen the genome for differentially methylated positions and regions; however, hits should be further validated using other methods. Manifold approaches exist to validate epigenome-wide findings or to perform candidate gene DNA methylation analysis. In recent years, the method of choice has been pyrosequencing, which is the most sensitive and accurate technology to quantify DNA methylation differences down to 2 %. However, pyrosequencing is rather laborious, expensive, not suitable for high-throughput, and can only analyze short DNA stretches including few CpG sites. Mass-spectrometry method like the Epityper is capable of analyzing up to 384 samples in parallel; longer DNA fragments (up to 600 bp) can be assayed for changes in methylation levels down to 5 %, but with less precision and imperfect resolution as not all CpGs in a given fragment can be analyzed. Cloning-based bisulfite sequencing or direct Sanger sequencing of bisulfite-specific PCR-products followed by Epigenetic Sequencing Methylation analysis (ESME) suffer from poor sensitivity (Lewin et al., 2004). In addition, they are very laborious, and are thus not suitable for high sample throughput. It can be expected that all these methods will soon be replaced by strategies using massive parallel- or deep sequencing – also called next-generation sequencing (NGS) – technology. This technological progress can also be used for locus-specific DNA methylation analysis by targeted deep bisulfite sequencing (Leitao et al., 2018), which tremendously reduces costs for high-throughput DNA methylation analysis, and offers outstanding quality in terms of sensitivity, accuracy and reproducibility.

Therefore, we outline a cost-efficient method, which allows for simultaneous analysis of longer genomic stretches at high resolution and appropriate sensitivity in larger cohorts that can be used to validate genome-wide data and to perform candidate gene approaches for many genes in parallel.

Based on a recently published protocol (Leitao et al., 2018), we present a PCR amplicon-based deep bisulfite sequencing approach for DNA methylation analysis of four candidate genes commonly assessed in psychoneuroendocrine research. This approach allows for simultaneous and affordable analysis of larger DNA-fragments (up to 400 bp) of multiple candidate genes at single base resolution in large cohorts in a single experiment.

To explore the validity of the assay, DNA methylation standards ranging from 0 % to 100 % for all fragments were assayed in triplicates. Reliability of the assays was tested from DNA of one female and one male donor, which was bisulfite treated in three independent trials and a pool generated out of the three conversions. Following, DNA from both donors was analyzed for reliability from three bisulfite converted DNAs and the pool in 10 separate PCR replicates each.

To illustrate the method, we investigated 90 female individuals (45 with a diagnosis of borderline personality disorder (BPD) and 45 matched healthy controls) for their DNA methylation pattern of genomic regions typically studied in stress research. Assays were performed for the *NR3C1 1F* promoter fragment including 42 CpGs (McGowan et al., 2009; Moser et al., 2007; Weaver et al., 2004), for the whole *SLC6A4* specific CpG island and 3 CpGs located upstream of the CpG island (84 CpGs; Alexander et al., 2014; Beach et al., 2010; Devlin et al., 2010; Kang et al., 2013; Philibert et al., 2007; Zhao et al., 2013), for five CpGs located in the genomic region coding for the *FKBP5* (Hohne et al., 2015; Klengel et al., 2013; Tyrka et al., 2015; Zannas et al., 2016), and for 12CpGs located in a putative enhancer element located in intron 3 of the *OXTR* (Gouin et al., 2017).

## Materials and Methods

### Principles of NGS library preparation

The NGS workflow for targeted bisulfite sequencing comprises four different steps: i) Sample preparation; ii) cluster generation; iii) sequencing; and iv) data analysis. As cluster generation and sequencing are part of the Illumina platform, and data analysis is already described elsewhere (Leitao et al., 2018; Rahmann, 2013) we will mainly focus on sample preparation here (see also Appendix 2).

For appropriate NGS library sample preparation, two rounds of polymerase chain reaction (PCR) are performed, starting from bisulfite converted DNA. In the first PCR, we used gene specific primers carrying tags for the second PCR (see Appendix 3). In the second PCR, specific adapter primers bind to these tag sequences and introduce additional DNA sequences to the PCR product, including barcode sequences. Each of these barcode sequences are unique to a specific NGS primer pair and serve as identifiers for a specific individual. In our study these NGS primer pairs consisted of eight different forward and 12 different reverse primers (based on Illumina adapters [N/S/E]501-[N/S/E]508 and N701-N712), which lead to 96 different combinations of primer pairs (as illustrated in Appendix 4). However, many more sets of primer pairs can be designed and used as identifiers for several hundreds of study participants (www.illumina.com).

To summarize, during first round PCR, all samples are amplified for a specific target region before the samples are re-amplified with specific adapter primers in a second PCR (Appendix 2). Subsequently, second round PCR products of each specific amplicon are pooled, purified by use of magnetic beads, and measured for their concentration. For sequencing, all pools of purified amplicons have to be adjusted for the same copy number to ensure that all samples are sequenced with the same probability.

### DNA Extraction, bisulfite conversion and PCRs

DNA was isolated from saliva samples using a salting-out procedure as described elsewhere (Miller et al., 1988). We used saliva for DNA extraction because of higher yields compared to buccal swab DNA. We deemed this necessary for the protocol establishment phase, and we are well aware that saliva-derived DNA does not represent the optimal source for epigenetic analyses as it includes a mixture of different cell-types (mucosa cells, leucocytes besides bacteria and leftovers).

Bisulfite conversion was performed from 500 ng genomic DNA using the *EZ DNA* Methylation-Gold Kit (Zymo Research, Irvine – USA) as it shows high conversion rates, integrity and amplifiability of DNA (Tierling et al., 2018).

One µl of bisulfite-converted DNA was subjected to PCR using the G2 Green Mix (Promega, Mannheim – Germany) with bisulfite-specific primers at temperatures and concentrations as indicated in Appendix 3. When using the Illumina MiSeq 2 × 250 v2 reagent kit, primers must be chosen to lead to maximum amplicon lengths of 400 bp. If possible, SNPs and CpG sites in the primer binding region should be avoided. If polymorphic regions in the primer target region cannot be avoided one may choose to introduce a degenerate site or “wobble” to compensate for potential variability in the target sequence. CpG sites in the primer binding sites should be kept to a minimum and are only allowed from the primers middle to the 5’ ends. If a potentially methylated cytosine could not be avoided, either degenerated primers or primers with a mismatch neither targeting the methylated nor the unmethylated sequence were used to prevent preferential allelic amplification. As DNA strands are no longer complementary after bisulfite conversion primers can be designed targeting the upper or lower strand. This leads to different orientations of sequencing directions, which is indicated by the arrows in Figure 1. All primers were tested by gradient PCR and different primer concentrations in order to achieve optimal conditions for second round PCRs. After quality control of first round PCR-products on a 1.5 % agarose gel (primer dimers and by-products must be strictly avoided), one µl of PCR products were subjected to second round PCR. After quality control on a 1.5 % agarose gel, four µl of each 2^nd^round PCR-product were pooled and purified using the Magnetic bead-based MagSi-NGS^*PREP*^ Plus Kit (MagnaMedics, Aachen; Germany) according to the manufacturers protocol. Subsequently, samples were quantified on a dual-channel fluorometer (Quantus TM Fluorometer, Promega, Mannheim; Germany), and adjusted for similar copy numbers for subsequent NGS analysis. The following paired-end sequencing was performed on a MiSeq system (Illumina, San Diego; USA) using the Illumina MiSeq reagent Kit v2 (500 cycles-2 × 250 paired end) in collaboration with the BioChip Labor of the Center of Medical Biotechnology (ZMB, University Duisburg/Essen).

**Figure 1:**
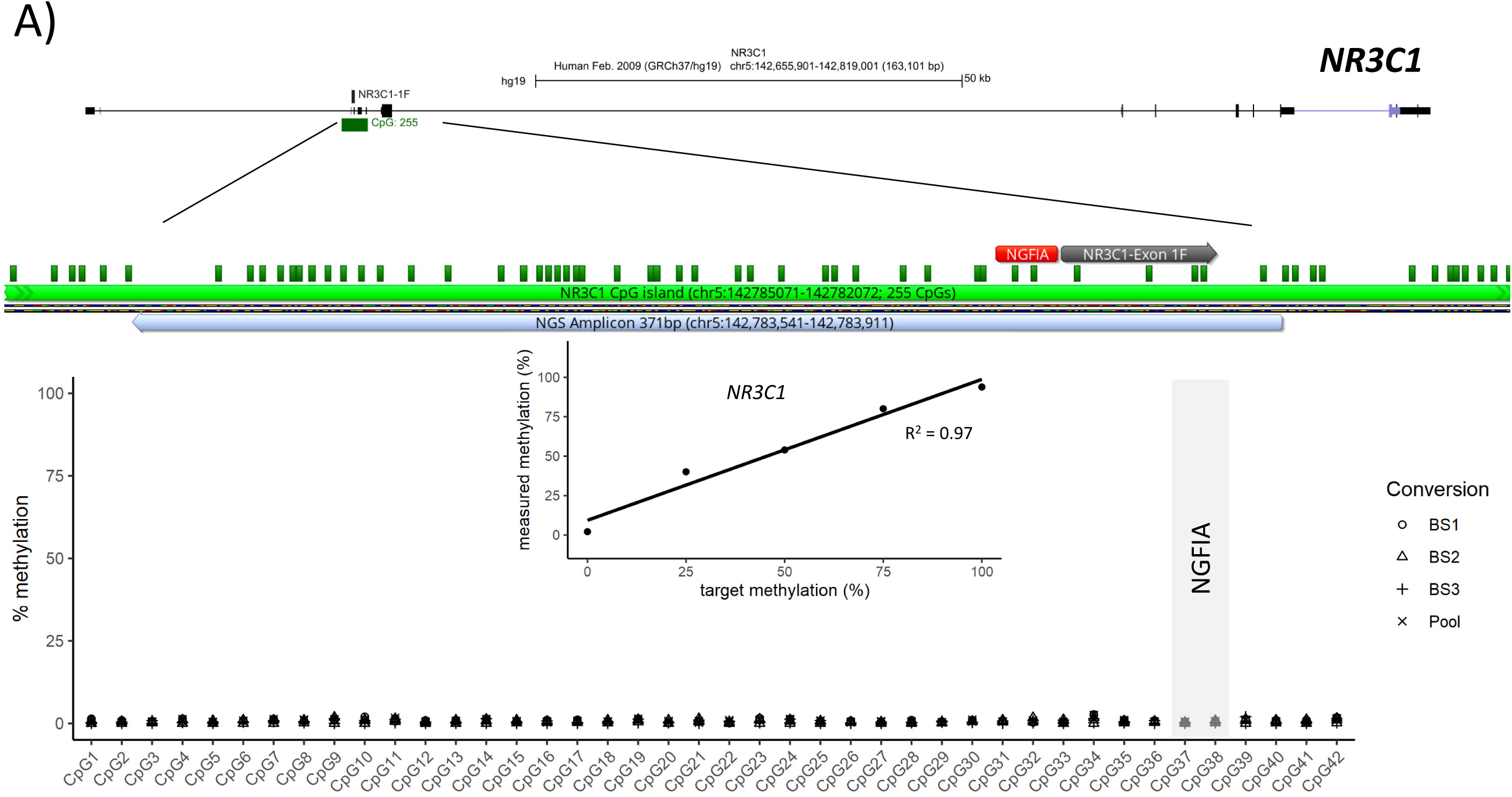
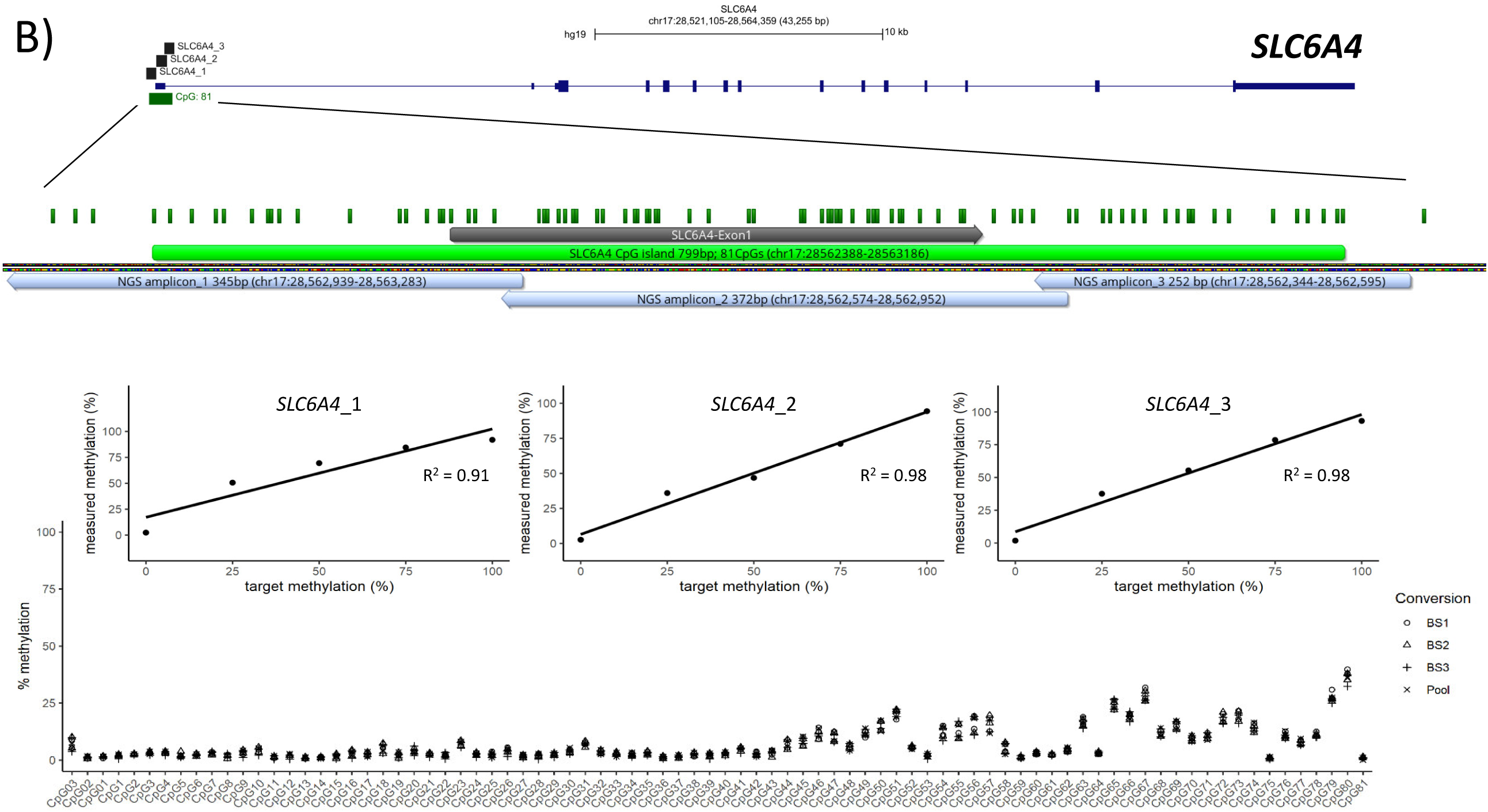
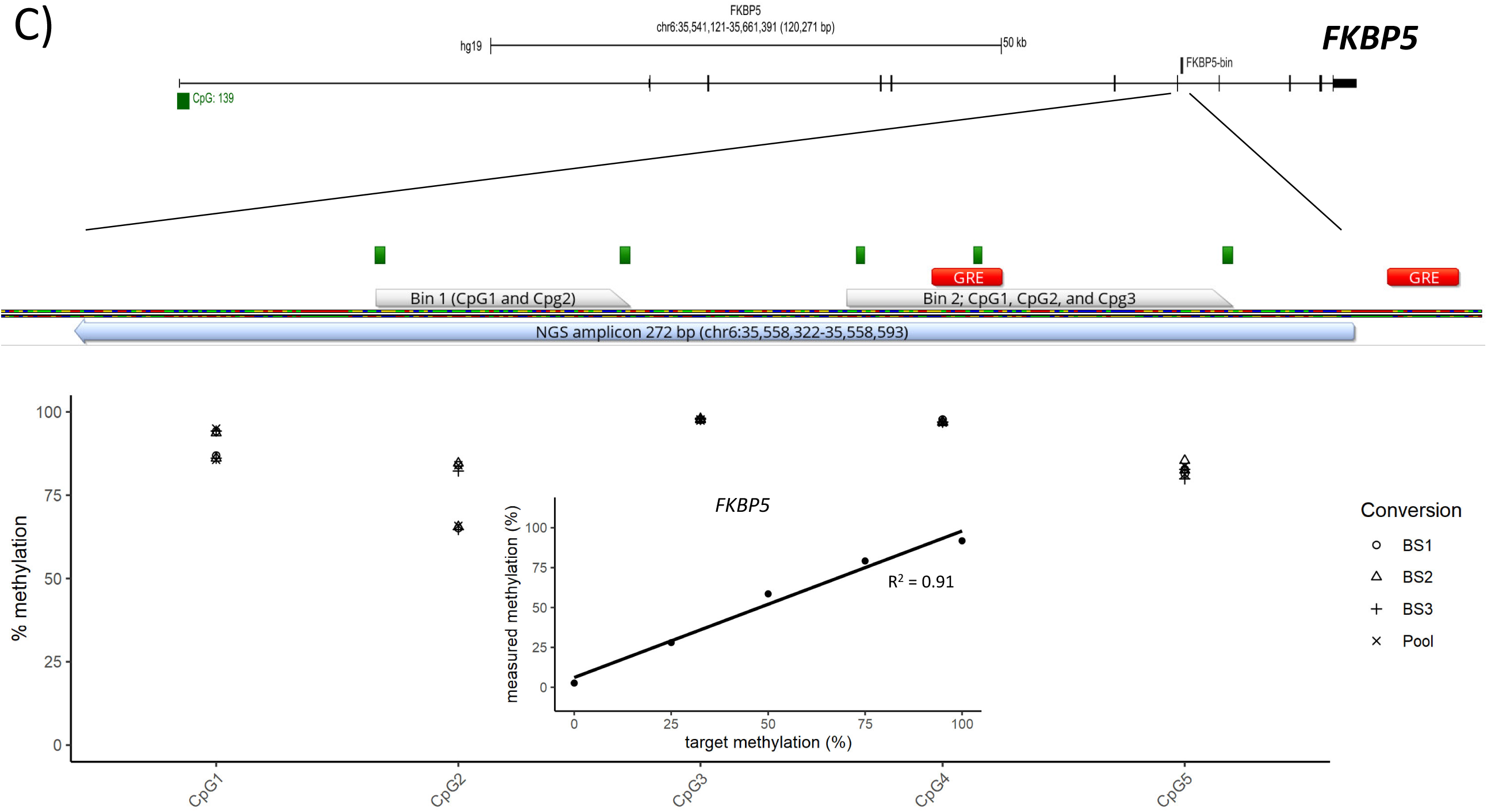
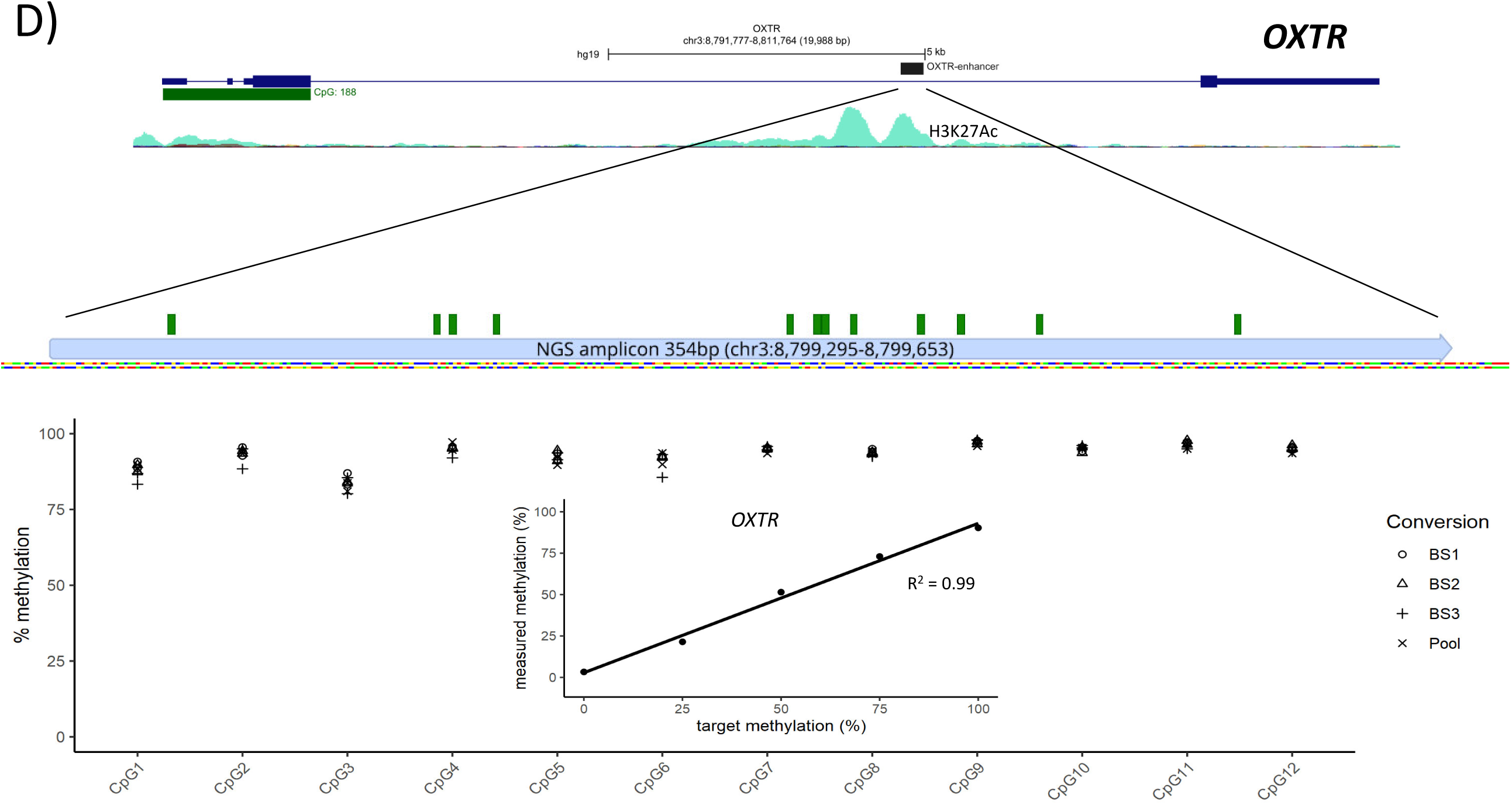
Chromosomal location of all amplicons and corresponding DNA methylation levels. Chromosomal position of all genes are illustrated using graphical outputs generated by the UCSC Genome Browser (https://genome.ucsc.edu) and Geneious Prime 2019 software (https://www.geneious.com). All genes are presented in 5’ -> 3’ orientation from left to right. Amplicons are highlighted with respect to their genomic orientation. All CpG sites are illustrated with green bars, CpG islands in light green, and exons in grey color. To determine the reliability of the assay, genomic DNA of a male and female donor were both bisulfite-treated in triplicates. From each triplicate and a pool of the triplicates 10 PCR replicates were prepared from all amplicons to test for the means standard deviations of the assays. Linearity of the assays is illustrated by plotting expected to experimental methylation values of a serial diluted methylation standard.

### Quality controls

To test the validity of the assays, five DNA methylation standards (100 %; 75 %; 50 %; 25 % and 0 % methylation) were also analyzed. Methylation standards (100 % and 0 %) were either prepared by *in vitro* methylation of human genomic DNA using M.SssI CpG methyltransferase (NEB, Frankfurt a.M; Germany) or by whole-genome amplification using the REPLI-g Mini Kit (Qiagen, Hilden; Germany). Methylation standards (75 %, 50 % and 25 %) were obtained by mixing 100 % and 0 % methylated DNA. For each amplicon methylation standards were assayed in triplicates. Linear regression was performed plotting the expected to measured DNA methylation values to calculate the coefficient of determination (R^2^). To determine the reliability of the assay, genomic DNA of a male and female donor were both bisulfite-treated in triplicates. From each triplicate and a pool of the triplicates 10 PCR replicates were prepared from all amplicons to test for the means standard deviations of the assays.

### Proof of principle

To demonstrate feasibility of the assay, DNA was taken from an existing cohort of female patients diagnosed with borderline personality disorder (BPD; n = 45) and 45 female healthy control participants (Flasbeck et al., 2018); a brief description of the cohort can be found in supplementary material 5. The study was approved by the local ethics committee of the Medical Faculty of the Ruhr University Bochum (project number 4639-13) and signed informed consent was obtained from all participants.

### Bioinformatics

For sequence analysis, FASTQ files from all sequencing reads were generated to enable third party software analysis using amplikyzer2 software (Leitao et al., 2018; Rahmann, 2013). This software enables to determine mean DNA methylation values of all CpG dinucleotides in a region, or of each CpG dinucleotide across all reads and allows comparing these values between different samples. Amplikyzer2 software was used with default settings for DNA methylation analysis excluding reads with less than 95 % bisulfite conversion rate and showing less coverage than 1000 reads per sample. In addition, we checked for the presence of SNPs, which might introduce or abolish CpG methylation sites. CpG sites created by SNPs were not included in the analysis. Unlike the standard setting, allele-specific DNA methylation was not calculated. Discrimination of allele specific methylation is only possible in the presence of heterozygous SNPs. These can be very helpful in imprinted regions and on the X-chromosome. As a further quality control, the amplikyzer2 graphical output was used to check for potential technical issues, which are indicated by bars of grey color; blue fields represent unmethylated CpGs and red color indicates methylated CpGs (Appendix 6).

### Statistical Analysis

Testing the validity of the assays, correlations between expected and observed percent DNA methylation of the DNA methylation standard curves were assessed by linear regression. Reliability of the assays was tested by the analysis of technical replicates. Mean standard deviation for all amplicons was calculated.

For a more comprehensive insight, we started with comparing the mean DNA methylation over all corresponding CpGs per analyzed fragment between patients with BPD and controls. The equality of variances was tested with Levene’s test, afterwards Student’s t-tests or Welch’s t-test were used respectively to compare the groups. In case of *SLC6A4*, where three fragments belong to the same candidate gene, differences in mean DNA methylation between the groups were analyzed separately with Student’s t-tests and adjusted for the multiple comparisons by Benjamini and Hochberg correction (Benjamini and Hochberg, 1995).

Further, we considered differences in DNA methylation levels between borderline patients and controls at single CpGs. Where Levene’s test indicated equal variances, Student’s t-tests were performed between the groups; otherwise Welch’s t-test was used. All p-values were adjusted by Benjamini and Hochberg correction for multiple testing by the number of CpGs tested for each candidate gene. We defined a P-value, or FDR-adjusted P-value, of <0.05 as statistically significant. The analyses were performed using R statistical programming (version 3.5.1).

## Results

Six different amplicons with a total length of 1973 basepairs, comprising 143 CpG sites were sequenced using the Illumina Miseq with the 2 × 250 v2 cycle kit. On average, assays showed around 7500 reads/sample (maximum 12025 and minimum 4591 reads per sample and amplicon). All samples were checked for sequencing quality using amplikyzer2 software generating alignment files and graphical outputs to identify regions of missing or unclear sequence information (Appendix 6). Unclear regions or CpG sites were not included in the further analysis.

Validity of the assays was first tested by the analysis of differentially methylated DNA standards (100 %, 75 %, 50 %, 25 % and 0 %), which revealed high linearity (R^2^> 0.9) for all amplicons (Figure 1). We also observed excellent reproducibility of results from two different DNAs – from a male and female donor-that were bisulfite-treated in three independent approaches and analyzed in 10 independent PCR reactions and a pool out of the three. An overview of DNA methylation for all genomic regions is illustrated in Figure 1; technical specifications of the assays are presented in Appendix 7.

### NR3C1

Forty-two CpG sites of the *NR3C1 1F* region as described elsewhere (McGowan et al., 2009) were investigated for their DNA methylation pattern. As indicated in the amplikyzer2-generated graphical output and data file, no technical problems occurred and all 42 CpGs located in our *NR3C1 1F* amplicon could be analyzed (Figure 2). Testing reliability of the assay, the DNA methylation analysis showed a mean standard deviation of 0.67 % (0.46 % for the tested female DNA; 0.88 % for the tested male DNA) with a maximum SD of 2.70 % and a high linearity (R^2^= 0.969) when analyzing methylation standards (Figure1; Appendix 7).

**Figure 2:**
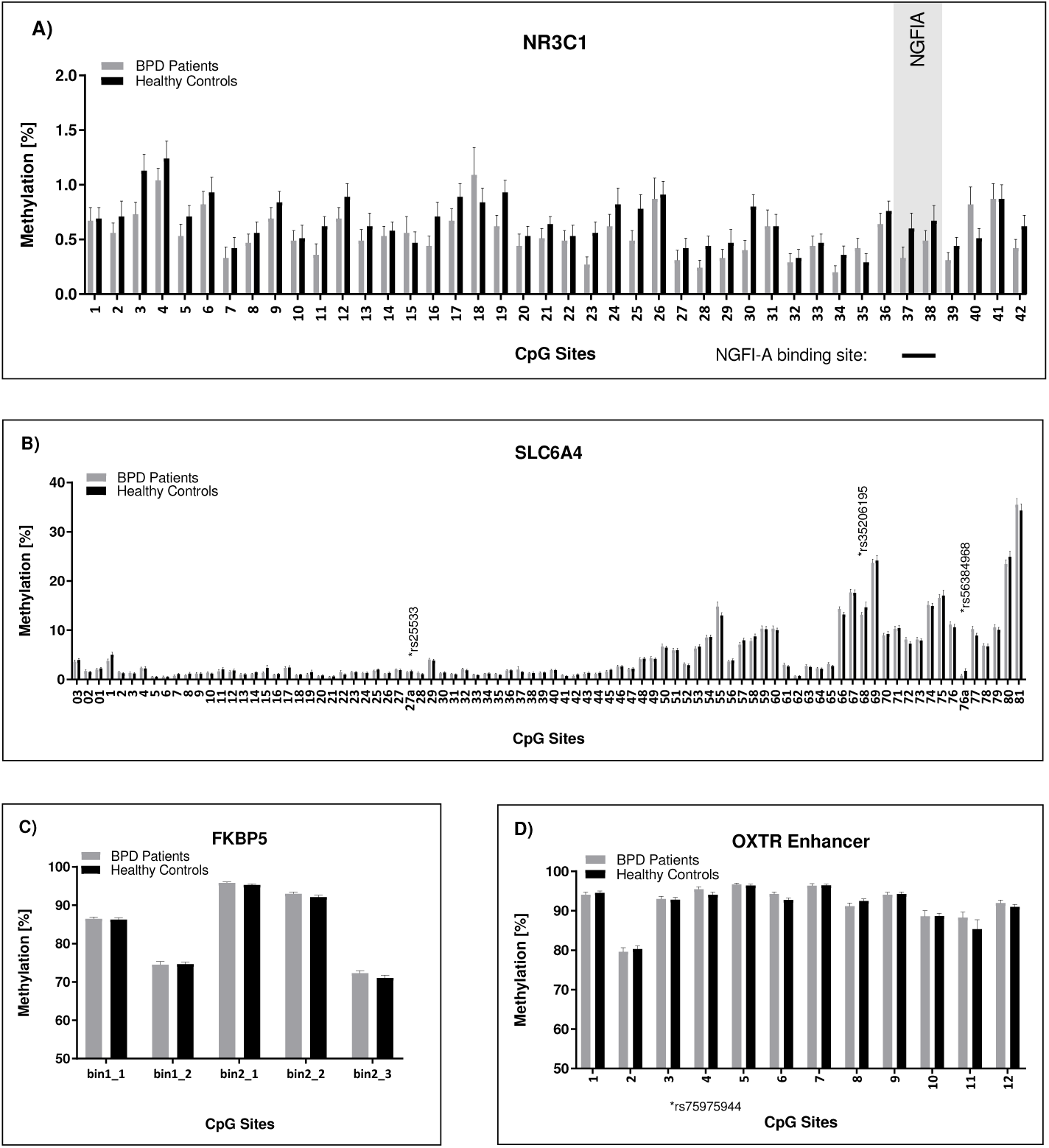
DNA methylation levels comparing BPD to healthy controls. Percentage methylation and standard deviation of all CpGs are presented for all amplicons/genes comparing BPD patients (grey) vs healthy controls (black)

We found low DNA methylation throughout all CpGs in all subjects (Figure 2). Borderline patients vs matched healthy controls were compared for their respective *NR3C1 1F* DNA methylation status, and as Levene’s test indicated unequal variances (F = 4.91, p = .029), degrees of freedom were adjusted from 88 to 80.84. Mean DNA methylation levels differed significantly by group according to a Welch’s t-test, t(80.84) = 3.01, p = .003. However, the difference between BPD patients and controls of 0.08 % were negligible (BPD: 0.66 % ± 0.02 %; control: 0.74 % ± 0.03 % (mean ± SEM)). The observed difference of 0.08 % cannot be reliably analyzed given the technical mean standard deviation of 0.46 % for this fragment.

### SLC6A4

The entire *SLC6A4* CpG island including 81 CpGs and 3 additional CpGs located upstream of the CpG island were investigated by the generation of three PCR amplicons. Genomic regions for rs25533 C/T (introduces a methylation site), rs35206195 C/T (disrupts a methylation site) and rs56384968 C/G (shifts the methylation site about one nucleotide) were not included in the statistical DNA methylation analysis. The methylation analysis for the three *SLC6A4* fragments showed a mean standard deviation between 1.37 % to 1.51 % with a maximum SD between 1.37 % to 6.1 % and a high linearity (R^2^> 0.9) when analyzing methylation standards (Figure 1 and Appendix 7).

Mean methylation of BPD patients and controls were identical (BPD: 4.76 % ± 0.14; control: 4.76 ± 0.12 % (mean ± SEM); Figure 2). No significant differences in DNA methylation between patients with BPD and controls in any of the three fragments (*SLC6A4*_1 t(88) = 1.76, FDR-adjusted p = .243; *SLC6A4*_2 t(88) = – 0.16, FDR-adjusted p = .873; *SLC6A4*_3 t(88) = −0.38, FDR-adjusted p = .873) were observed.

### FKBP5

A 272 base pair fragment of a *FKBP5* intron 7 specific fragment, containing five CpGs of the bin 1 and bin 2 fragments (Klengel et al., 2013) was also subjected to targeted deep bisulfite sequencing analysis. The assay showed high linearity (R^2^= 0.98) in the identification of differentially methylated standards and high reproducibility (mean SD: 1.29 %) when analyzing 10 independent PCR products generated from 2 subjects from 3 independent bisulfite modified DNAs (Figure 1; Appendix 7). We could not find a significant difference in mean methylation between BPD patients and controls (Figure 2; BPD: 84.43 % ± 0.37; control: 83.90 ± 0.36 % (mean ± SEM)); (t(88) = −1.02, p = .311).

### OXTR

We initially focused on 12 CpGs located 3 kb downstream of rs53576 in a putative enhancer fragment of *OXTR* intron 3, a region that has been associated with empathy, loneliness, more sensitive parenting techniques, and with lower rates of autism (Kim et al., 2010). Polymorphic sites at rs114435510, rs75975944, and rs7636061 where handled with special caution in which only rs7636061 C/T abolishes a methylation site but cannot be differentiated in bisulfite converted DNA. The assays showed high linearity (R^2^> 0.99) and a mean standard deviation of 2.24 % (Figure 1 and Appendix 7). We found an almost fully methylated enhancer region in all investigated participants (BPD: 91.69 % ± 0.40 %; control: 91.41 % ± 0.28 % (mean ± SEM)). Comparing DNA methylation from patients with BPD to healthy controls, no significant differences could be observed (t(88) = −0.524, p = .602; Figure 2).

## Discussion

Here, we illustrate a new experimental protocol for cost efficient high-throughput DNA methylation analysis in four different genes of relevance for stress research. We first demonstrated excellent reliability and validity of the new NGS-based approach (Figure 1, 2 and Appendix 7). As a proof of principle, we investigated percentage DNA methylation of 142 CpGs located in six different amplicons representing four different genes of saliva-derived DNA from 45 female BPD patients compared to matched healthy controls. The whole procedure was performed according to a recently published, highly flexible protocol (Leitao et al., 2018) that can either be used for the investigation of large cohorts for a limited number of genomic regions or to investigate smaller cohorts for a larger amount of genetic elements.

Chen et al. (2017) were the first in the field of psychiatric research to describe the application of targeted bisulfite sequencing for accurate detection of 5-methylcytosine and also 5-hydroxmethylcytosine. Using a strategy applying three independent PCRs they investigated 16 different post-mortem human brain tissues with greater than 100 x coverage. Targeting three different regions of the Opioid Receptor Kappa 1 (*OPRK1*) gene containing 83 CpG sites they could confirm high precision of their assay (R^2^> 0.75) when compared to results obtained by RRBS. Comparing their assay to the assay presented here, our assay is less laborious as it needs only two rounds of PCRs followed by one final DNA purification step. In addition, every amplicon pool is adjusted for copy numbers, which are inserted into the sequencing reaction leading to predictable high coverages (Appendix 2). As illustrated, our assays show high linearity, and reproducibility for the six investigated amplicons (Figure 1, 2 and Appendix 7). These data are highly similar to recently published work introducing a comparable NGS approach for the DNA methylation analysis of the *FKBP5* gene (Roeh et al., 2018). Performing ***H****igh* ***A****ccurate* ***M****ethod for* ***T****argeted* ***B****isulfite* ***S****equencing* (HAM-TBS) a mean sensitivity of 0.72 % was reported, compared to a mean sensitivity between 0.67 % to 2.24 % we achieved for our six different assays (see Appendix 7). Whether the slightly higher mean standard deviation identified for our assay is due to its technical properties or due to the heterogeneous tissue used for DNA extraction has to be further evaluated in the future.

As illustrated in supplemental figure 8, *FKBP5* revealed specific differences at CpG site 2 of Bin 1. Whether this an inter-individual variation or gender specific variation in the DNA-methylation signature cannot be clarified here. However, inter-individual differences in DNA methylation and the larger sample size assayed here could have led also to a slightly higher mean standard deviation comparing our assay to the HAM-TBS approach. However, for both platforms this is a much higher sensitivity compared to those achieved by pyrosequencing or the Epityper. Compared to our method, HAM-TBS requires only one PCR and thus is less susceptible to cross-contamination and PCR biases. Nevertheless, as the second PCR in our approach introduces barcode labelled primers this enables enormous flexibility for multiplexing of samples and amplicons. Around 15 million usable reads can be generated during one NGS run, which means that 15.000 different amplicons with 1000 reads per sample can be successfully investigated per run. In addition, this higher throughput leads to cost reduction depending on the amount of different amplicons that are finally assayed. Compared to other methods like the Epityper, pyrosequencing and to traditional Sanger sequencing, the method presented here is much cheaper, more accurate, less laborious, and allows for higher sample throughput (Appendix 1).

For *NR3C1*, we present an assay to analyze DNA methylation of the promoter of exon 1F, most commonly investigated when differential DNA methylation of *NR3C1* is of interest. As also observed by others (Elwenspoek et al., 2019; McGowan et al., 2009; Moser et al., 2007), the analyzed *NR3C1* stretch was generally unmethylated, with mean DNA methylation values below 1 %. The statistical difference for *NR3C1* we observed comparing individuals with BPD and controls has to be handled with caution. First, given that the sensitivity of our method for this gene fragment is 0.46 %, differences in DNA methylation of 0.08 % cannot be reliably assessed. Second, DNA was extracted from saliva, which represents a heterogeneous tissue, and introduces a potential bias into the analysis (Jones et al., 2018; Kumsta, 2019). Third, individual genotype influencing individual DNA methylation signatures cannot be excluded (Gertz et al., 2011; Schroder et al., 2017). Fourth, the investigated cohort was rather small (Heijmans and Mill, 2012) and last but not least, differences in DNA methylation of way below 1 % are unlikely to be of biological significance (Heijmans and Mill, 2012; Matthijs et al., 2016).

For *SLC6A4*, we were able to capture the entire CpG island, divided into three NGS amplicons. Comparing borderline patients vs healthy controls we found almost identical values in the whole *SLC6A4* CpG island. Interestingly, the method described here provides the possibility to achieve a visual impression of whole CpG island methylation. In case of *SLC6A4* the whole CpG island is rather unmethylated, with a slight increase in DNA methylation beginning at the end of exon 1 and a further increase downstream of the CpG island (Figure 2). This finding is supported by literature describing that methylation of CpG shores, defined as the 2 kb of sequence flanking a CpG island, tend to be more dynamic than methylation of the CpG island itself (Bird, 2002; Greally, 2013).

For *FKBP5*, we focused on 5 CpGs in intron 7 of the *FKBP5* gene, two of which are located in a glucocorticoid receptor consensus binding site. A previous study has reported that DNA methylation of so-called bin 2 (encompassing 3 CpGs sites) was associated with childhood adversity in a genotype-dependent manner (Klengel et al., 2013), which makes this region a promising candidate for the investigation of *gene* by *environment* by *DNA methylation* interaction. We observed no differences between BPD and controls. Interestingly, significant inter-individual differences were observed comparing female and male DNA methylation signatures for this gene fragment. This underlines the necessity to control for inter-individual and gender-mediated differences when performing DNA methylation analysis (Bock et al., 2008; Boks et al., 2009).

The investigation of the whole *OXTR* specific CpG island with 188 CpGs present in a stretch of 2318 nucleotides would have been desirable here, but the enormous density of CpGs and redundancy of DNA after bisulfite treatment make specific primer design at this genomic locus extremely challenging. Focusing on a putative enhancer element located in the *OXTR* intron 3, we found high DNA methylation levels, but no differences between groups. Interestingly, as already described by others, gene bodies and intronic sequences show high methylation levels whereas CpG islands tend to be rarely methylated (Eckhardt et al., 2006; Zhou et al., 2011).

## Summary

When designing and performing targeted deep bisulfite sequencing assays, the following aspects should be taken into account:

In order to prevent technical issues, SNPs and CpG sites in the primer binding region should be avoided or should be faced with strategies as illustrated above. Furthermore, any formation of primer dimers and unspecific by-products must be avoided. This makes primer design and efficient generation of high quality first round PCR-products very difficult when facing large CpG islands – like in cases of the *OXTR -* and demands for thorough primer design and careful optimization and validation of reliable assays. In addition critical care must be taken to prevent cross contamination when transferring first round PCR-products to prepare second round PCR.

Analyzing DNAs from a male and a female donor, which were bisulfite-treated in three independent assays and a pool generated from theses samples revealed almost identical results with the exception of *FKBP5*, where inter-individual differences in the DNA methylation profile were visible. As the conversion rate of most commercial bisulfite assays is above 99 % (Tierling et al., 2018) in our opinion it is sufficient to perform bisulfite treatments in unique copies. In addition, the amplikyzer2 software enables filtering of insufficient converted DNAs for final analysis. Analysis of pooled bisulfite converted DNA as suggested by others (Roeh et al., 2018) might complicate the analysis as according to our experience one bisulfite converted DNA of lower quality negatively affects the quality of the whole pool and leads to results of lower quality. Thus, we would recommend performing single bisulfite-treatment and to perform PCRs in triplicates, as long as sufficient bisulfite-treated DNA is available. In cases of low DNA availability, for instance when using buccal swabs for cell collection, performing bisulfite-treatment and PCRs in unique copies should also be acceptable.

As suggested by others (Heijmans and Mill, 2012), we recommend to always use homogenous cells for DNA methylation analysis. Analysis of mouthwash and buccal cell DNA is often complicated by fluctuating amounts of blood-cell derived DNA, which influences epigenetic results. Blood consists of multiple different cell types with their respective divergent epigenotypes. As blood cell composition can vary across time and is influenced by several factors, including inflammation, physical and/or psychosocial stress, sexual hormones etc., many differences in DNA methylation might be driven by altered blood cell composition and not by cell type-specific alterations of DNA methylation (Hummel et al., 2018; Jones et al., 2018; Kumsta, 2019). When using blood, this challenge can partly be addressed via physical isolation of cells via cell sorting or immunomagnetic isolation (Schwaiger et al., 2016), or by using blood cell counts to strip away variations in DNA methylation that can be attributed to differences in leukocyte subset composition (Jones et al., 2018).

## Conclusion

The method we outlined here can be used for targeted analysis of selected candidate genes and for validation of hits derived from whole-genome or array-based approaches. It represents a cost-efficient, highly sensitive method that offers increased spatial resolution compared to other methods. Furthermore, it enables high-throughput analysis and offers the possibility of parallel investigation of many candidate genes in the same run. We expect this method to contribute to greatly improved quality and quantity of DNA methylation data and thus to improved understanding of the association between DNA methylation variation in different phenotypes and diseases.

## References

Alexander, N., Wankerl, M., Hennig, J., Miller, R., Zankert, S., Steudte-Schmiedgen, S., Stalder, T., Kirschbaum, C., 2014. DNA methylation profiles within the serotonin transporter gene moderate the association of 5-HTTLPR and cortisol stress reactivity. Transl Psychiat 4.

Argentieri, M.A., Nagarajan, S., Seddighzadeh, B., Baccarelli, A.A., Shields, A.E., 2017. Epigenetic Pathways in Human Disease: The Impact of DNA Methylation on Stress-Related Pathogenesis and Current Challenges in Biomarker Development. Ebiomedicine 18, 327–350.

Beach, S.R.H., Brody, G.H., Todorov, A.A., Gunter, T.D., Philibert, R.A., 2010. Methylation at SLC6A4 Is Linked to Family History of Child Abuse: An Examination of the Iowa Adoptee Sample. Am J Med Genet B 153b, 710–713.

Benjamini, Y., Hochberg, Y., 1995. Controlling the False Discovery Rate – a Practical and Powerful Approach to Multiple Testing. J R Stat Soc B 57, 289–300.

Bibikova, M., Barnes, B., Tsan, C., Ho, V., Klotzle, B., Le, J.M., Delano, D., Zhang, L., Schroth, G.P., Gunderson, K.L., Fan, J.B., Shen, R., 2011. High density DNA methylation array with single CpG site resolution. Genomics 98, 288–295.

Bird, A., 2002. DNA methylation patterns and epigenetic memory. Gene Dev 16, 6–21.

Bock, C., Walter, J., Paulsen, M., Lengauer, T., 2008. Inter-individual variation of DNA methylation and its implications for large-scale epigenome mapping. Nucleic Acids Res 36, e55.

Boks, M.P., Derks, E.M., Weisenberger, D.J., Strengman, E., Janson, E., Sommer, I.E., Kahn, R.S., Ophoff, R.A., 2009. The relationship of DNA methylation with age, gender and genotype in twins and healthy controls. Plos One 4, e6767.

Cecil, C.A., Lysenko, L.J., Jaffee, S.R., Pingault, J.B., Smith, R.G., Relton, C.L., Woodward, G., McArdle, W., Mill, J., Barker, E.D., 2014. Environmental risk, Oxytocin Receptor Gene (OXTR) methylation and youth callous-unemotional traits: a 13-year longitudinal study. Mol Psychiatry 19, 1071–1077.

Chen, G.G., Gross, J.A., Lutz, P.E., Vaillancourt, K., Maussion, G., Bramoulle, A., Theroux, J.F., Gardini, E.S., Ehlert, U., Bourret, G., Masurel, A., Lepage, P., Mechawar, N., Turecki, G., Ernst, C., 2017. Medium throughput bisulfite sequencing for accurate detection of 5-methylcytosine and 5-hydroxymethylcytosine. BMC Genomics 18, 96.

Cokus, S.J., Feng, S., Zhang, X., Chen, Z., Merriman, B., Haudenschild, C.D., Pradhan, S., Nelson, S.F., Pellegrini, M., Jacobsen, S.E., 2008. Shotgun bisulphite sequencing of the Arabidopsis genome reveals DNA methylation patterning. Nature 452, 215–219.

Dammann, G., Teschler, S., Haag, T., Altmuller, F., Tuczek, F., Dammann, R.H., 2011. Increased DNA methylation of neuropsychiatric genes occurs in borderline personality disorder. Epigenetics 6, 1454–1462.

Devlin, A.M., Brain, U., Austin, J., Oberlander, T.F., 2010. Prenatal exposure to maternal depressed mood and the MTHFR C677T variant affect SLC6A4 methylation in infants at birth. PLoS One 5, e12201.

Eckhardt, F., Lewin, J., Cortese, R., Rakyan, V.K., Attwood, J., Burger, M., Burton, J., Cox, T.V., Davies, R., Down, T.A., Haefliger, C., Horton, R., Howe, K., Jackson, D.K., Kunde, J., Koenig, C., Liddle, J., Niblett, D., Otto, T., Pettett, R., Seemann, S., Thompson, C., West, T., Rogers, J., Olek, A., Berlin, K., Beck, S., 2006. DNA methylation profiling of human chromosomes 6, 20 and 22. Nat Genet 38, 1378–1385.

Edwards, J.R., O’Donnell, A.H., Rollins, R.A., Peckham, H.E., Lee, C., Milekic, M.H., Chanrion, B., Fu, Y., Su, T., Hibshoosh, H., Gingrich, J.A., Haghighi, F., Nutter, R., Bestor, T.H., 2010. Chromatin and sequence features that define the fine and gross structure of genomic methylation patterns. Genome Res 20, 972–980.

Elwenspoek, M.M.C., Hengesch, X., Leenen, F.A.D., Sias, K., Fernandes, S.B., Schaan, V.K., Meriaux, S.B., Schmitz, S., Bonnemberger, F., Schachinger, H., Vogele, C., Muller, C.P., Turner, J.D., 2019. Glucocorticoid receptor signaling in leukocytes after early life adversity. Dev Psychopathol, 1–11.

Feil, R., Fraga, M.F., 2012. Epigenetics and the environment: emerging patterns and implications. Nat Rev Genet 13, 97–109.

Flasbeck, V., Moser, D., Kumsta, R., Brune, M., 2018. The OXTR Single-Nucleotide Polymorphism rs53576 Moderates the Impact of Childhood Maltreatment on Empathy for Social Pain in Female Participants: Evidence for Differential Susceptibility. Front Psychiatry 9.

Gertz, J., Varley, K.E., Reddy, T.E., Bowling, K.M., Pauli, F., Parker, S.L., Kucera, K.S., Willard, H.F., Myers, R.M., 2011. Analysis of DNA methylation in a three-generation family reveals widespread genetic influence on epigenetic regulation. PLoS Genet 7, e1002228.

Godfrey, K.M., Costello, P.M., Lillycrop, K.A., 2015. The developmental environment, epigenetic biomarkers and long-term health. J Dev Orig Hlth Dis 6, 399–406.

Gouin, J.P., Zhou, Q.Q., Booij, L., Boivin, M., Cote, S.M., Hebert, M., Ouellet-Morin, I., Szyf, M., Tremblay, R.E., Turecki, G., Vitaro, F., 2017. Associations among oxytocin receptor gene (OXTR) DNA methylation in adulthood, exposure to early life adversity, and childhood trajectories of anxiousness. Sci Rep-Uk 7.

Greally, J.M., 2013. DNA METHYLATION Bidding the CpG island goodbye. Elife 2.

Heijmans, B.T., Mill, J., 2012. Commentary: The seven plagues of epigenetic epidemiology. Int J Epidemiol 41, 74–78.

Hohne, N., Poidinger, M., Merz, F., Pfister, H., Bruckl, T., Zimmermann, P., Uhr, M., Holsboer, F., Ising, M., 2015. FKBP5 Genotype-Dependent DNA Methylation and mRNA Regulation After Psychosocial Stress in Remitted Depression and Healthy Controls. Int J Neuropsychoph 18.

Hummel, E.M., Hessas, E., Muller, S., Beiter, T., Fisch, M., Eibl, A., Wolf, O.T., Giebel, B., Platen, P., Kumsta, R., Moser, D.A., 2018. Cell-free DNA release under psychosocial and physical stress conditions. Transl Psychiat 8.

Jones, M.J., Moore, S.R., Kobor, M.S., 2018. Principles and Challenges of Applying Epigenetic Epidemiology to Psychology. Annu Rev Psychol 69, 459–485.

Kang, H.J., Kim, J.M., Stewart, R., Kim, S.Y., Bae, K.Y., Kim, S.W., Shin, I.S., Shin, M.G., Yoon, J.S., 2013. Association of SLC6A4 methylation with early adversity, characteristics and outcomes in depression. Prog Neuro-Psychoph 44, 23–28.

Kim, H.S., Sherman, D.K., Sasaki, J.Y., Xu, J., Chu, T.Q., Ryu, C., Suh, E.M., Graham, K., Taylor, S.E., 2010. Culture, distress, and oxytocin receptor polymorphism (OXTR) interact to influence emotional support seeking. P Natl Acad Sci USA 107, 15717–15721.

Klengel, T., Mehta, D., Anacker, C., Rex-Haffner, M., Pruessner, J.C., Pariante, C.M., Pace, T.W.W., Mercer, K.B., Mayberg, H.S., Bradley, B., Nemeroff, C.B., Holsboer, F., Heim, C.M., Ressler, K.J., Rein, T., Binder, E.B., 2013. Allele-specific FKBP5 DNA demethylation mediates gene-childhood trauma interactions. Nat Neurosci 16, 33–41.

Kumsta, R., 2019. The role of epigenetics for understanding mental health difficulties and its implications for psychotherapy research. Psychol Psychother-T 92, 190–207.

Laurent, L., Wong, E., Li, G., Huynh, T., Tsirigos, A., Ong, C.T., Low, H.M., Sung, K.W.K., Rigoutsos, I., Loring, J., Wei, C.L., 2010. Dynamic changes in the human methylome during differentiation. Genome Research 20, 320–331.

Leitao, E., Beygo, J., Zeschnigk, M., Klein-Hitpass, L., Bargull, M., Rahmann, S., Horsthemke, B., 2018. Locus-Specific DNA Methylation Analysis by Targeted Deep Bisulfite Sequencing. Methods Mol Biol 1767, 351–366.

Lewin, J., Schmitt, A.O., Adorjan, P., Hildmann, T., Piepenbrock, C., 2004. Quantitative DNA methylation analysis based on four-dye trace data from direct sequencing of PCR amplificates. Bioinformatics 20, 3005–3012.

Lister, R., Pelizzola, M., Dowen, R.H., Hawkins, R.D., Hon, G., Tonti-Filippini, J., Nery, J.R., Lee, L., Ye, Z., Ngo, Q.M., Edsall, L., Antosiewicz-Bourget, J., Stewart, R., Ruotti, V., Millar, A.H., Thomson, J.A., Ren, B., Ecker, J.R., 2009. Human DNA methylomes at base resolution show widespread epigenomic differences. Nature 462, 315–322.

Matthijs, G., Souche, E., Alders, M., Corveleyn, A., Eck, S., Feenstra, I., Race, V., Sistermans, E., Sturm, M., Weiss, M., Yntema, H., Bakker, E., Scheffer, H., Bauer, P., 2016. Guidelines for diagnostic next-generation sequencing. Eur J Hum Genet 24, 2–5.

McGowan, P.O., Sasaki, A., D’Alessio, A.C., Dymov, S., Labonte, B., Szyf, M., Turecki, G., Meaney, M.J., 2009. Epigenetic regulation of the glucocorticoid receptor in human brain associates with childhood abuse. Nat Neurosci 12, 342–348.

Meissner, A., Gnirke, A., Bell, G.W., Ramsahoye, B., Lander, E.S., Jaenisch, R., 2005. Reduced representation bisulfite sequencing for comparative high-resolution DNA methylation analysis. Nucleic Acids Research 33, 5868–5877.

Miller, S.A., Dykes, D.D., Polesky, H.F., 1988. A Simple Salting out Procedure for Extracting DNA from Human Nucleated Cells. Nucleic Acids Research 16, 1215–1215.

Moser, D., Molitor, A., Kumsta, R., Tatschner, T., Riederer, P., Meyer, J., 2007. The glucocorticoid receptor gene exon 1-F promoter is not methylated at the NGFI-A binding site in human hippocampus. World J Biol Psychiatry 8, 262–268.

Philibert, R., Madan, A., Andersen, A., Cadoret, R., Packer, H., Sandhu, H., 2007. Serotonin transporter mRNA levels are associated with the methylation of an upstream CpG island. Am J Med Genet B Neuropsychiatr Genet 144B, 101–105.

Pidsley, R., Zotenko, E., Peters, T.J., Lawrence, M.G., Risbridger, G.P., Molloy, P., Van Djik, S., Muhlhausler, B., Stirzaker, C., Clark, S.J., 2016. Critical evaluation of the Illumina MethylationEPIC BeadChip microarray for whole-genome DNA methylation profiling. Genome Biol 17, 208.

Rahmann, S., Beygo, J., Kanber, D., Martin, M.N., Horsthemke, B., Buiting, K.,, 2013. Amplikyzer: Automated methylation analysis of amplicons from bisulfite flowgram sequencing. PeerJ PrePrints.

Roeh, S., Wiechmann, T., Sauer, S., Kodel, M., Binder, E.B., Provencal, N., 2018. HAM-TBS: high-accuracy methylation measurements via targeted bisulfite sequencing. Epigenet Chromatin 11.

Schroder, C., Leitao, E., Wallner, S., Schmitz, G., Klein-Hitpass, L., Sinha, A., Jockel, K.H., Heilmann-Heimbach, S., Hoffmann, P., Nothen, M.M., Steffens, M., Ebert, P., Rahmann, S., Horsthemke, B., 2017. Regions of common inter-individual DNA methylation differences in human monocytes: genetic basis and potential function. Epigenetics Chromatin 10, 37.

Schwaiger, M., Grinberg, M., Moser, D., Zang, J.C.S., Heinrichs, M., Hengstler, J.G., Rahnenfuhrer, J., Cole, S., Kumsta, R., 2016. Altered Stress-Induced Regulation of Genes in Monocytes in Adults with a History of Childhood Adversity. Neuropsychopharmacol 41, 2530–2540.

Tierling, S., Schmitt, B., Walter, J., 2018. Comprehensive Evaluation of Commercial Bisulfite-Based DNA Methylation Kits and Development of an Alternative Protocol With Improved Conversion Performance. Genet Epigenetics 10.

Turecki, G., Meaney, M.J., 2016. Effects of the Social Environment and Stress on Glucocorticoid Receptor Gene Methylation: A Systematic Review. Biol Psychiatry 79, 87–96.

Tyrka, A.R., Ridout, K.K., Parade, S.H., Paquette, A., Marsit, C.J., Seifer, R., 2015. Childhood maltreatment and methylation of FK506 binding protein 5 gene (FKBP5). Dev Psychopathol 27, 1637–1645.

Unternaehrer, E., Luers, P., Mill, J., Dempster, E., Meyer, A.H., Staehli, S., Lieb, R., Hellhammer, D.H., Meinlschmidt, G., 2012. Dynamic changes in DNA methylation of stress-associated genes (OXTR, BDNF) after acute psychosocial stress. Transl Psychiatry 2, e150.

Urich, M.A., Nery, J.R., Lister, R., Schmitz, R.J., Ecker, J.R., 2015. MethylC-seq library preparation for base-resolution whole-genome bisulfite sequencing. Nat Protoc 10, 475–483.

van Ijzendoorn, M.H., Caspers, K., Bakermans-Kranenburg, M.J., Beach, S.R.H., Philibert, R., 2010. Methylation Matters: Interaction Between Methylation Density and Serotonin Transporter Genotype Predicts Unresolved Loss or Trauma. Biol Psychiat 68, 405–407.

Weaver, I.C.G., Cervoni, N., Champagne, F.A., D’Alessio, A.C., Sharma, S., Seckl, J.R., Dymov, S., Szyf, M., Meaney, M.J., 2004. Epigenetic programming by maternal behavior. Nat Neurosci 7, 847–854.

Zannas, A.S., Wiechmann, T., Gassen, N.C., Binder, E.B., 2016. Gene-Stress-Epigenetic Regulation of FKBP5: Clinical and Translational Implications. Neuropsychopharmacology 41, 261–274.

Zhao, J.Y., Goldberg, J., Bremner, J.D., Vaccarino, V., 2013. Association Between Promoter Methylation of Serotonin Transporter Gene and Depressive Symptoms: A Monozygotic Twin Study. Psychosom Med 75, 523–529.

Zhou, V.W., Goren, A., Bernstein, B.E., 2011. Charting histone modifications and the functional organization of mammalian genomes. Nature Reviews Genetics 12, 7–18.

