## Appendix 1-8 for "Targeted bisulfite sequencing: A novel tool for the assessment of DNA methylation with high sensitivity and increased coverage"

**Appendix 1: Comparison of DNA methylation platforms**

| | Sequence length | coverage | genotyping | throughput | Cost/sample | sensitivity | bioinformatics |
|---|---|---|---|---|---|---|---|
| **Targeted bisulfite sequencing** | up to 400 bp | full | yes | up to 15.000 samples per run* | Very low | best | required |
| **Ham-TBS** | up to 400 bp | full | yes | up to 96 samples per run | low | best | required |
| **Pyrosequencer** | 50 - 100 bp | full | yes | up to 96 samples per run | moderate | very good 2% - 5% | not required |
| **Epityper** | 200 - 600 bp | incomplete | no | up to 384 samples per run | moderate | good ~5% | not required |
| **Cloning-based Sanger sequencing** | 1000 bp | full | yes | up to 96 samples per run | moderate | moderate | not required |
| **Direct Sanger sequencing-ESME** | 1000 bp | full | yes | up to 96 samples per run | moderate | very low 10% - 20% | required |

* based on 1000 reads/sample on average

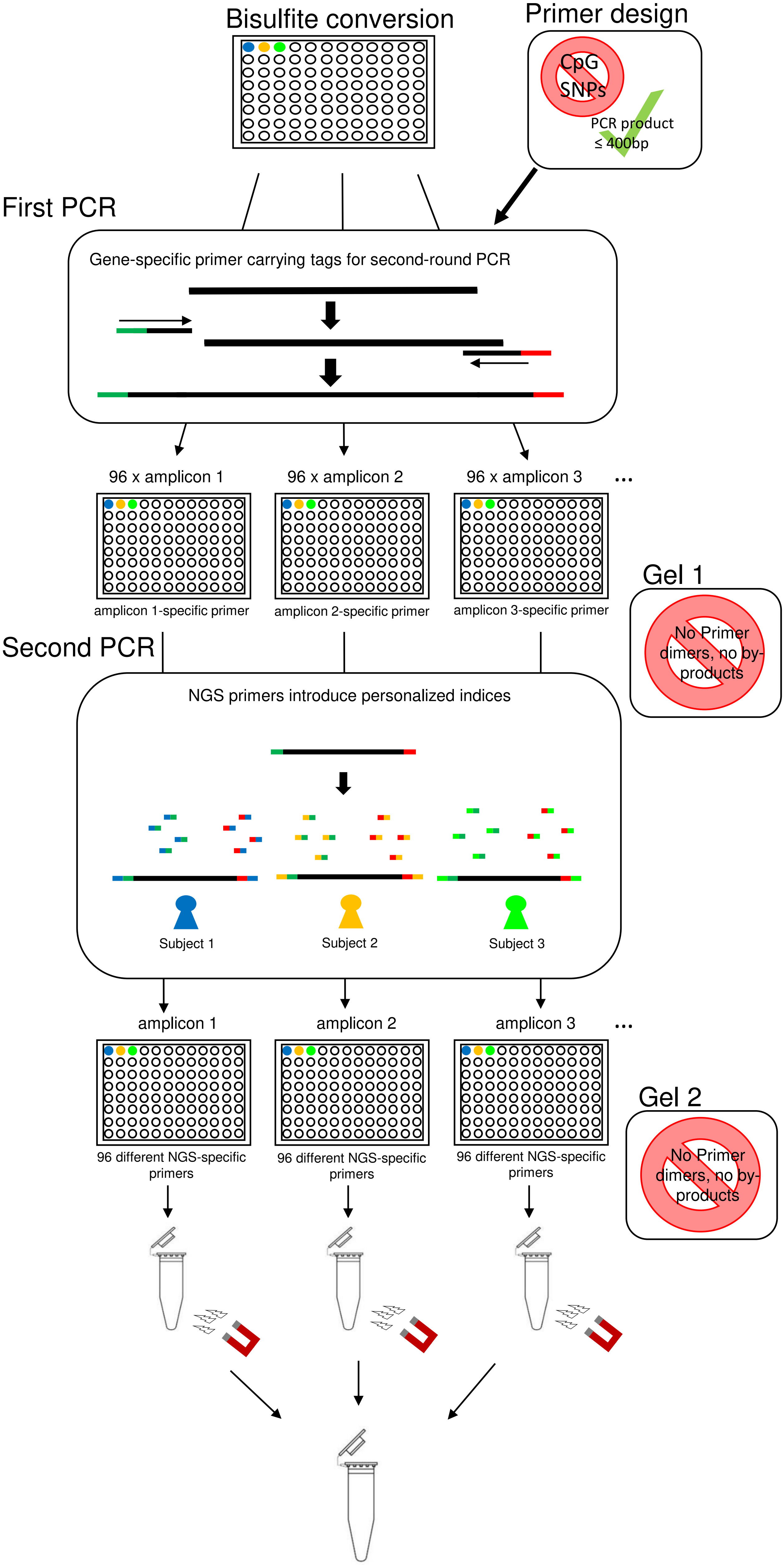
Bisulfite conversion
Primer design
CpG
SNPs
PCR product
≤ 400bp
First PCR
Gene-specific primer carrying tags for second-round PCR
96 x amplicon 1
96 x amplicon 2
96 x amplicon 3
...
amplicon 1-specific primer
amplicon 2-specific primer
amplicon 3-specific primer
Gel 1
No Primer dimers, no by-products
Second PCR
NGS primers introduce personalized indices
Subject 1
Subject 2
Subject 3
amplicon 1
amplicon 2
amplicon 3
...
Gel 2
No Primer dimers, no by-products
96 different NGS-specific primers
96 different NGS-specific primers
96 different NGS-specific primers

**Appendix 2: Workflow for targeted bisulfite sequencing**

For appropriate NGS library sample preparation, two rounds of polymerase chain reaction (PCR) are performed, starting from bisulfite converted DNA. In the first PCR, gene specific primers carrying tags for the second PCR are used. In the second PCR, specific adapter primers bind to these tag sequences and introduce additional DNA sequences to the PCR product, including barcode sequences. Each of these barcode sequences are unique to a specific NGS primer pair and serve as identifiers for a specific individual. Finally, all gene-specific PCR products of n subjects (here 96) are purified by use of magnetic beads to be sequenced together.

**Appendix 3: Technical and experimental amplicon characteristics**

| Name | Sequences forward (Fw) and reverse (Rv) primers | Chromosomal location (bp) | Primer concentrations/annealing | Number of CpGs |
|---|---|---|---|---|
| ***NR3C1 1F*** | Fw-CTTGCTTCCTGGCACGAGgggggtagatttggtttttt<br>Rv-CAGGAAACAGCTATGACtcccttccctaaaacctc | chr5:142,783,541-142,783,911 (371) | 0.1µM/56.9°C | 42 |
| ***SLC6A4_1*** | Fw-CTTGCTTCCTGGCACGAGtaggaggggagggatttt<br>Rv-CAGGAAACAGCTATGACaaacctctaaactaMactcacatc | chr17:28,562,939-28,563,283 (345) | 0.2µM/59.4°C | 23 |
| ***SLC6A4_2*** | Fw-CTTGCTTCCTGGCACGAGgggaagaaggtttggaaaga<br>Rv-CAGGAAACAGCTATGACtccctcccctcctaactctaa | chr17:28,562,574-28,562,952 (379) | 0.2µM/59.4°C | 42 |
| ***SLC6A4_3*** | Fw-CTTGCTTCCTGGCACGAGagaggttgtaaagtttttgtaataatagat<br>Rv-CAGGAAACAGCTATGACtttctttccaaaccttcttcc | chr17:28,562,344-28,562,595 (252) | 0.15µM/59.8°C | 19 |
| ***FKBP5*** | Fw-CTTGCTTCCTGGCACGAGttttgggttgaggatagaaagg<br>Rv-CAGGAAACAGCTATGACatccaaaacaactaacaaattctct | chr6:35,558,322-35,558,593 (272) | 0.1µM/56°C | 5 |
| ***OXTR_Enh*** | Fw-CTTGCTTCCTGGCACGAGttgtgggtaggagtaggatttta<br>Rv-CAGGAAACAGCTATGACttctcatcctaaatctaaaaatcactt | chr3:8,799,262-8,799,615 (354) | 0.15µM/59.4°C | 12 |

Forward (Fw) und reverse (Rv) primers are displayed for six different amplicons. Capital letters represent tag sequences, lower case represent bisulfite-specific gene sequences. Chromosomal position, number of CpGs and PCR conditions for the respective amplicons are depicted.

**Appendix 4: NGS primers**

| Forward Primers | | Flow cell binding sequence | Index (i5) | Sequencing primer binding site | Variable sequence | Tag sequence |
|---|---|---|---|---|---|---|
| 1 | [N/S/E]501 | AATGATACGGCGACCACCGAGATCTACAC | TAGATCGC | ACACTCTTTCCCTACACGACGCTCTTCCGATCT | TATAGC | CTTGCTTCCTGGCACGAG |
| 2 | [N/S/E]502 | AATGATACGGCGACCACCGAGATCTACAC | CTCTCTAT | ACACTCTTTCCCTACACGACGCTCTTCCGATCT | ATAGAG | CTTGCTTCCTGGCACGAG |
| 3 | [N/S/E]503 | AATGATACGGCGACCACCGAGATCTACAC | TATCCTCT | ACACTCTTTCCCTACACGACGCTCTTCCGATCT | CCTATC | CTTGCTTCCTGGCACGAG |
| 4 | [N/S/E]504 | AATGATACGGCGACCACCGAGATCTACAC | AGAGTAGA | ACACTCTTTCCCTACACGACGCTCTTCCGATCT | GGCTCT | CTTGCTTCCTGGCACGAG |
| 5 | [N/S/E]505 | AATGATACGGCGACCACCGAGATCTACAC | GTAAGGAG | ACACTCTTTCCCTACACGACGCTCTTCCGATCT | AGGCGA | CTTGCTTCCTGGCACGAG |
| 6 | [N/S/E]506 | AATGATACGGCGACCACCGAGATCTACAC | ACTGCATA | ACACTCTTTCCCTACACGACGCTCTTCCGATCT | TAATCT | CTTGCTTCCTGGCACGAG |
| 7 | [N/S/E]507 | AATGATACGGCGACCACCGAGATCTACAC | AAGGAGTA | ACACTCTTTCCCTACACGACGCTCTTCCGATCT | CAGGAC | CTTGCTTCCTGGCACGAG |
| 8 | [N/S/E]508 | AATGATACGGCGACCACCGAGATCTACAC | CTAAGCCT | ACACTCTTTCCCTACACGACGCTCTTCCGATCT | GTACTG | CTTGCTTCCTGGCACGAG |

| Reverse Primers | | Flow cell binding sequence | Index (i7) | Sequencing primer binding site | Tag sequence |
|---|---|---|---|---|---|
| 1 | N701 | CAAGCAGAAGACGGCATACGAGAT | TCGCCTTA | GTGACTGGAGTTCAGACGTGTGCTCTTCCGATCT | CAGGAAACAGCTATGAC |
| 2 | N702 | CAAGCAGAAGACGGCATACGAGAT | CTAGTACG | GTGACTGGAGTTCAGACGTGTGCTCTTCCGATCT | CAGGAAACAGCTATGAC |
| 3 | N703 | CAAGCAGAAGACGGCATACGAGAT | TTCTGCCT | GTGACTGGAGTTCAGACGTGTGCTCTTCCGATCT | CAGGAAACAGCTATGAC |
| 4 | N704 | CAAGCAGAAGACGGCATACGAGAT | GCTCAGGA | GTGACTGGAGTTCAGACGTGTGCTCTTCCGATCT | CAGGAAACAGCTATGAC |
| 5 | N705 | CAAGCAGAAGACGGCATACGAGAT | AGGAGTCC | GTGACTGGAGTTCAGACGTGTGCTCTTCCGATCT | CAGGAAACAGCTATGAC |
| 6 | N706 | CAAGCAGAAGACGGCATACGAGAT | CATGCCTA | GTGACTGGAGTTCAGACGTGTGCTCTTCCGATCT | CAGGAAACAGCTATGAC |
| 7 | N707 | CAAGCAGAAGACGGCATACGAGAT | GTAGAGAG | GTGACTGGAGTTCAGACGTGTGCTCTTCCGATCT | CAGGAAACAGCTATGAC |
| 8 | N708 | CAAGCAGAAGACGGCATACGAGAT | CCTCTCTG | GTGACTGGAGTTCAGACGTGTGCTCTTCCGATCT | CAGGAAACAGCTATGAC |
| 9 | N709 | CAAGCAGAAGACGGCATACGAGAT | AGCGTAGC | GTGACTGGAGTTCAGACGTGTGCTCTTCCGATCT | CAGGAAACAGCTATGAC |
| 10 | N710 | CAAGCAGAAGACGGCATACGAGAT | CAGCCTCG | GTGACTGGAGTTCAGACGTGTGCTCTTCCGATCT | CAGGAAACAGCTATGAC |
| 11 | N711 | CAAGCAGAAGACGGCATACGAGAT | TGCCTCTT | GTGACTGGAGTTCAGACGTGTGCTCTTCCGATCT | CAGGAAACAGCTATGAC |
| 12 | N712 | CAAGCAGAAGACGGCATACGAGAT | TCCTCTAC | GTGACTGGAGTTCAGACGTGTGCTCTTCCGATCT | CAGGAAACAGCTATGAC |

| Number | Name | Flow cell binding sequence | Index (i5) | Sequencing primer binding site | Variable sequence | Tag sequence | Name | Flow cell binding sequence | Index (i7) | Sequencing primer binding site | Tag sequence |
|---|---|---|---|---|---|---|---|---|---|---|---|
| NGS #1 | [N/S/E]501 | AATGATACGGCGACCACCGAGATCTACAC | TAGATCGC | ACACTCTTTCCCTACACGACGCTCTTCCGATCT | TATAGC | CTTGCTTCCTGGCACGAG | N701 | CAAGCAGAAGACGGCATACGAGAT | TCGCCTTA | GTGACTGGAGTTCAGACGTGTGCTCTTCCGATCT | CAGGAAACAGCTATGAC |
| NGS #2 | [N/S/E]501 | AATGATACGGCGACCACCGAGATCTACAC | TAGATCGC | ACACTCTTTCCCTACACGACGCTCTTCCGATCT | TATAGC | CTTGCTTCCTGGCACGAG | N702 | CAAGCAGAAGACGGCATACGAGAT | CTAGTACG | GTGACTGGAGTTCAGACGTGTGCTCTTCCGATCT | CAGGAAACAGCTATGAC |
| NGS #3 | [N/S/E]501 | AATGATACGGCGACCACCGAGATCTACAC | TAGATCGC | ACACTCTTTCCCTACACGACGCTCTTCCGATCT | TATAGC | CTTGCTTCCTGGCACGAG | N703 | CAAGCAGAAGACGGCATACGAGAT | TTCTGCCT | GTGACTGGAGTTCAGACGTGTGCTCTTCCGATCT | CAGGAAACAGCTATGAC |
| NGS #4 | [N/S/E]501 | AATGATACGGCGACCACCGAGATCTACAC | TAGATCGC | ACACTCTTTCCCTACACGACGCTCTTCCGATCT | TATAGC | CTTGCTTCCTGGCACGAG | N704 | CAAGCAGAAGACGGCATACGAGAT | GCTCAGGA | GTGACTGGAGTTCAGACGTGTGCTCTTCCGATCT | CAGGAAACAGCTATGAC |
| NGS #5 | [N/S/E]501 | AATGATACGGCGACCACCGAGATCTACAC | TAGATCGC | ACACTCTTTCCCTACACGACGCTCTTCCGATCT | TATAGC | CTTGCTTCCTGGCACGAG | N705 | CAAGCAGAAGACGGCATACGAGAT | AGGAGTCC | GTGACTGGAGTTCAGACGTGTGCTCTTCCGATCT | CAGGAAACAGCTATGAC |
| NGS #6 | [N/S/E]501 | AATGATACGGCGACCACCGAGATCTACAC | TAGATCGC | ACACTCTTTCCCTACACGACGCTCTTCCGATCT | TATAGC | CTTGCTTCCTGGCACGAG | N706 | CAAGCAGAAGACGGCATACGAGAT | CATGCCTA | GTGACTGGAGTTCAGACGTGTGCTCTTCCGATCT | CAGGAAACAGCTATGAC |
| NGS #7 | [N/S/E]501 | AATGATACGGCGACCACCGAGATCTACAC | TAGATCGC | ACACTCTTTCCCTACACGACGCTCTTCCGATCT | TATAGC | CTTGCTTCCTGGCACGAG | N707 | CAAGCAGAAGACGGCATACGAGAT | GTAGAGAG | GTGACTGGAGTTCAGACGTGTGCTCTTCCGATCT | CAGGAAACAGCTATGAC |
| NGS #8 | [N/S/E]501 | AATGATACGGCGACCACCGAGATCTACAC | TAGATCGC | ACACTCTTTCCCTACACGACGCTCTTCCGATCT | TATAGC | CTTGCTTCCTGGCACGAG | N708 | CAAGCAGAAGACGGCATACGAGAT | CCTCTCTG | GTGACTGGAGTTCAGACGTGTGCTCTTCCGATCT | CAGGAAACAGCTATGAC |
| NGS #9 | [N/S/E]501 | AATGATACGGCGACCACCGAGATCTACAC | TAGATCGC | ACACTCTTTCCCTACACGACGCTCTTCCGATCT | TATAGC | CTTGCTTCCTGGCACGAG | N709 | CAAGCAGAAGACGGCATACGAGAT | AGCGTAGC | GTGACTGGAGTTCAGACGTGTGCTCTTCCGATCT | CAGGAAACAGCTATGAC |
| NGS #10 | [N/S/E]501 | AATGATACGGCGACCACCGAGATCTACAC | TAGATCGC | ACACTCTTTCCCTACACGACGCTCTTCCGATCT | TATAGC | CTTGCTTCCTGGCACGAG | N710 | CAAGCAGAAGACGGCATACGAGAT | CAGCCTCG | GTGACTGGAGTTCAGACGTGTGCTCTTCCGATCT | CAGGAAACAGCTATGAC |
| NGS #11 | [N/S/E]501 | AATGATACGGCGACCACCGAGATCTACAC | TAGATCGC | ACACTCTTTCCCTACACGACGCTCTTCCGATCT | TATAGC | CTTGCTTCCTGGCACGAG | N711 | CAAGCAGAAGACGGCATACGAGAT | TGCCTCTT | GTGACTGGAGTTCAGACGTGTGCTCTTCCGATCT | CAGGAAACAGCTATGAC |
| NGS #12 | [N/S/E]501 | AATGATACGGCGACCACCGAGATCTACAC | TAGATCGC | ACACTCTTTCCCTACACGACGCTCTTCCGATCT | TATAGC | CTTGCTTCCTGGCACGAG | N712 | CAAGCAGAAGACGGCATACGAGAT | TCCTCTAC | GTGACTGGAGTTCAGACGTGTGCTCTTCCGATCT | CAGGAAACAGCTATGAC |
| NGS #13 | [N/S/E]502 | AATGATACGGCGACCACCGAGATCTACAC | CTCTCTAT | ACACTCTTTCCCTACACGACGCTCTTCCGATCT | ATAGAG | CTTGCTTCCTGGCACGAG | N701 | CAAGCAGAAGACGGCATACGAGAT | TCGCCTTA | GTGACTGGAGTTCAGACGTGTGCTCTTCCGATCT | CAGGAAACAGCTATGAC |
| NGS #14 | [N/S/E]502 | AATGATACGGCGACCACCGAGATCTACAC | CTCTCTAT | ACACTCTTTCCCTACACGACGCTCTTCCGATCT | ATAGAG | CTTGCTTCCTGGCACGAG | N702 | CAAGCAGAAGACGGCATACGAGAT | CTAGTACG | GTGACTGGAGTTCAGACGTGTGCTCTTCCGATCT | CAGGAAACAGCTATGAC |
| NGS #15 | [N/S/E]502 | AATGATACGGCGACCACCGAGATCTACAC | CTCTCTAT | ACACTCTTTCCCTACACGACGCTCTTCCGATCT | ATAGAG | CTTGCTTCCTGGCACGAG | N703 | CAAGCAGAAGACGGCATACGAGAT | TTCTGCCT | GTGACTGGAGTTCAGACGTGTGCTCTTCCGATCT | CAGGAAACAGCTATGAC |
| NGS #16 | [N/S/E]502 | AATGATACGGCGACCACCGAGATCTACAC | CTCTCTAT | ACACTCTTTCCCTACACGACGCTCTTCCGATCT | ATAGAG | CTTGCTTCCTGGCACGAG | N704 | CAAGCAGAAGACGGCATACGAGAT | GCTCAGGA | GTGACTGGAGTTCAGACGTGTGCTCTTCCGATCT | CAGGAAACAGCTATGAC |
| NGS #17 | [N/S/E]502 | AATGATACGGCGACCACCGAGATCTACAC | CTCTCTAT | ACACTCTTTCCCTACACGACGCTCTTCCGATCT | ATAGAG | CTTGCTTCCTGGCACGAG | N705 | CAAGCAGAAGACGGCATACGAGAT | AGGAGTCC | GTGACTGGAGTTCAGACGTGTGCTCTTCCGATCT | CAGGAAACAGCTATGAC |
| NGS #18 | [N/S/E]502 | AATGATACGGCGACCACCGAGATCTACAC | CTCTCTAT | ACACTCTTTCCCTACACGACGCTCTTCCGATCT | ATAGAG | CTTGCTTCCTGGCACGAG | N706 | CAAGCAGAAGACGGCATACGAGAT | CATGCCTA | GTGACTGGAGTTCAGACGTGTGCTCTTCCGATCT | CAGGAAACAGCTATGAC |
| NGS #19 | [N/S/E]502 | AATGATACGGCGACCACCGAGATCTACAC | CTCTCTAT | ACACTCTTTCCCTACACGACGCTCTTCCGATCT | ATAGAG | CTTGCTTCCTGGCACGAG | N707 | CAAGCAGAAGACGGCATACGAGAT | GTAGAGAG | GTGACTGGAGTTCAGACGTGTGCTCTTCCGATCT | CAGGAAACAGCTATGAC |
| NGS #20 | [N/S/E]502 | AATGATACGGCGACCACCGAGATCTACAC | CTCTCTAT | ACACTCTTTCCCTACACGACGCTCTTCCGATCT | ATAGAG | CTTGCTTCCTGGCACGAG | N708 | CAAGCAGAAGACGGCATACGAGAT | CCTCTCTG | GTGACTGGAGTTCAGACGTGTGCTCTTCCGATCT | CAGGAAACAGCTATGAC |
| NGS #21 | [N/S/E]502 | AATGATACGGCGACCACCGAGATCTACAC | CTCTCTAT | ACACTCTTTCCCTACACGACGCTCTTCCGATCT | ATAGAG | CTTGCTTCCTGGCACGAG | N709 | CAAGCAGAAGACGGCATACGAGAT | AGCGTAGC | GTGACTGGAGTTCAGACGTGTGCTCTTCCGATCT | CAGGAAACAGCTATGAC |
| NGS #22 | [N/S/E]502 | AATGATACGGCGACCACCGAGATCTACAC | CTCTCTAT | ACACTCTTTCCCTACACGACGCTCTTCCGATCT | ATAGAG | CTTGCTTCCTGGCACGAG | N710 | CAAGCAGAAGACGGCATACGAGAT | CAGCCTCG | GTGACTGGAGTTCAGACGTGTGCTCTTCCGATCT | CAGGAAACAGCTATGAC |
| NGS #23 | [N/S/E]502 | AATGATACGGCGACCACCGAGATCTACAC | CTCTCTAT | ACACTCTTTCCCTACACGACGCTCTTCCGATCT | ATAGAG | CTTGCTTCCTGGCACGAG | N711 | CAAGCAGAAGACGGCATACGAGAT | TGCCTCTT | GTGACTGGAGTTCAGACGTGTGCTCTTCCGATCT | CAGGAAACAGCTATGAC |
| NGS #24 | [N/S/E]502 | AATGATACGGCGACCACCGAGATCTACAC | CTCTCTAT | ACACTCTTTCCCTACACGACGCTCTTCCGATCT | ATAGAG | CTTGCTTCCTGGCACGAG | N712 | CAAGCAGAAGACGGCATACGAGAT | TCCTCTAC | GTGACTGGAGTTCAGACGTGTGCTCTTCCGATCT | CAGGAAACAGCTATGAC |
| NGS #25 | [N/S/E]503 | AATGATACGGCGACCACCGAGATCTACAC | TATCCTCT | ACACTCTTTCCCTACACGACGCTCTTCCGATCT | CCTATC | CTTGCTTCCTGGCACGAG | N701 | CAAGCAGAAGACGGCATACGAGAT | TCGCCTTA | GTGACTGGAGTTCAGACGTGTGCTCTTCCGATCT | CAGGAAACAGCTATGAC |
| NGS #26 | [N/S/E]503 | AATGATACGGCGACCACCGAGATCTACAC | TATCCTCT | ACACTCTTTCCCTACACGACGCTCTTCCGATCT | CCTATC | CTTGCTTCCTGGCACGAG | N702 | CAAGCAGAAGACGGCATACGAGAT | CTAGTACG | GTGACTGGAGTTCAGACGTGTGCTCTTCCGATCT | CAGGAAACAGCTATGAC |
| NGS #27 | [N/S/E]503 | AATGATACGGCGACCACCGAGATCTACAC | TATCCTCT | ACACTCTTTCCCTACACGACGCTCTTCCGATCT | CCTATC | CTTGCTTCCTGGCACGAG | N703 | CAAGCAGAAGACGGCATACGAGAT | TTCTGCCT | GTGACTGGAGTTCAGACGTGTGCTCTTCCGATCT | CAGGAAACAGCTATGAC |
| NGS #28 | [N/S/E]503 | AATGATACGGCGACCACCGAGATCTACAC | TATCCTCT | ACACTCTTTCCCTACACGACGCTCTTCCGATCT | CCTATC | CTTGCTTCCTGGCACGAG | N704 | CAAGCAGAAGACGGCATACGAGAT | GCTCAGGA | GTGACTGGAGTTCAGACGTGTGCTCTTCCGATCT | CAGGAAACAGCTATGAC |
| NGS #29 | [N/S/E]503 | AATGATACGGCGACCACCGAGATCTACAC | TATCCTCT | ACACTCTTTCCCTACACGACGCTCTTCCGATCT | CCTATC | CTTGCTTCCTGGCACGAG | N705 | CAAGCAGAAGACGGCATACGAGAT | AGGAGTCC | GTGACTGGAGTTCAGACGTGTGCTCTTCCGATCT | CAGGAAACAGCTATGAC |
| NGS #30 | [N/S/E]503 | AATGATACGGCGACCACCGAGATCTACAC | TATCCTCT | ACACTCTTTCCCTACACGACGCTCTTCCGATCT | CCTATC | CTTGCTTCCTGGCACGAG | N706 | CAAGCAGAAGACGGCATACGAGAT | CATGCCTA | GTGACTGGAGTTCAGACGTGTGCTCTTCCGATCT | CAGGAAACAGCTATGAC |
| NGS #31 | [N/S/E]503 | AATGATACGGCGACCACCGAGATCTACAC | TATCCTCT | ACACTCTTTCCCTACACGACGCTCTTCCGATCT | CCTATC | CTTGCTTCCTGGCACGAG | N707 | CAAGCAGAAGACGGCATACGAGAT | GTAGAGAG | GTGACTGGAGTTCAGACGTGTGCTCTTCCGATCT | CAGGAAACAGCTATGAC |
| NGS #32 | [N/S/E]503 | AATGATACGGCGACCACCGAGATCTACAC | TATCCTCT | ACACTCTTTCCCTACACGACGCTCTTCCGATCT | CCTATC | CTTGCTTCCTGGCACGAG | N708 | CAAGCAGAAGACGGCATACGAGAT | CCTCTCTG | GTGACTGGAGTTCAGACGTGTGCTCTTCCGATCT | CAGGAAACAGCTATGAC |
| NGS #33 | [N/S/E]503 | AATGATACGGCGACCACCGAGATCTACAC | TATCCTCT | ACACTCTTTCCCTACACGACGCTCTTCCGATCT | CCTATC | CTTGCTTCCTGGCACGAG | N709 | CAAGCAGAAGACGGCATACGAGAT | AGCGTAGC | GTGACTGGAGTTCAGACGTGTGCTCTTCCGATCT | CAGGAAACAGCTATGAC |
| NGS #34 | [N/S/E]503 | AATGATACGGCGACCACCGAGATCTACAC | TATCCTCT | ACACTCTTTCCCTACACGACGCTCTTCCGATCT | CCTATC | CTTGCTTCCTGGCACGAG | N710 | CAAGCAGAAGACGGCATACGAGAT | CAGCCTCG | GTGACTGGAGTTCAGACGTGTGCTCTTCCGATCT | CAGGAAACAGCTATGAC |
| NGS #35 | [N/S/E]503 | AATGATACGGCGACCACCGAGATCTACAC | TATCCTCT | ACACTCTTTCCCTACACGACGCTCTTCCGATCT | CCTATC | CTTGCTTCCTGGCACGAG | N711 | CAAGCAGAAGACGGCATACGAGAT | TGCCTCTT | GTGACTGGAGTTCAGACGTGTGCTCTTCCGATCT | CAGGAAACAGCTATGAC |
| NGS #36 | [N/S/E]503 | AATGATACGGCGACCACCGAGATCTACAC | TATCCTCT | ACACTCTTTCCCTACACGACGCTCTTCCGATCT | CCTATC | CTTGCTTCCTGGCACGAG | N712 | CAAGCAGAAGACGGCATACGAGAT | TCCTCTAC | GTGACTGGAGTTCAGACGTGTGCTCTTCCGATCT | CAGGAAACAGCTATGAC |
| NGS #37 | [N/S/E]504 | AATGATACGGCGACCACCGAGATCTACAC | AGAGTAGA | ACACTCTTTCCCTACACGACGCTCTTCCGATCT | GGCTCT | CTTGCTTCCTGGCACGAG | N701 | CAAGCAGAAGACGGCATACGAGAT | TCGCCTTA | GTGACTGGAGTTCAGACGTGTGCTCTTCCGATCT | CAGGAAACAGCTATGAC |
| NGS #38 | [N/S/E]504 | AATGATACGGCGACCACCGAGATCTACAC | AGAGTAGA | ACACTCTTTCCCTACACGACGCTCTTCCGATCT | GGCTCT | CTTGCTTCCTGGCACGAG | N702 | CAAGCAGAAGACGGCATACGAGAT | CTAGTACG | GTGACTGGAGTTCAGACGTGTGCTCTTCCGATCT | CAGGAAACAGCTATGAC |
| NGS #39 | [N/S/E]504 | AATGATACGGCGACCACCGAGATCTACAC | AGAGTAGA | ACACTCTTTCCCTACACGACGCTCTTCCGATCT | GGCTCT | CTTGCTTCCTGGCACGAG | N703 | CAAGCAGAAGACGGCATACGAGAT | TTCTGCCT | GTGACTGGAGTTCAGACGTGTGCTCTTCCGATCT | CAGGAAACAGCTATGAC |
| NGS #40 | [N/S/E]504 | AATGATACGGCGACCACCGAGATCTACAC | AGAGTAGA | ACACTCTTTCCCTACACGACGCTCTTCCGATCT | GGCTCT | CTTGCTTCCTGGCACGAG | N704 | CAAGCAGAAGACGGCATACGAGAT | GCTCAGGA | GTGACTGGAGTTCAGACGTGTGCTCTTCCGATCT | CAGGAAACAGCTATGAC |
| NGS #41 | [N/S/E]504 | AATGATACGGCGACCACCGAGATCTACAC | AGAGTAGA | ACACTCTTTCCCTACACGACGCTCTTCCGATCT | GGCTCT | CTTGCTTCCTGGCACGAG | N705 | CAAGCAGAAGACGGCATACGAGAT | AGGAGTCC | GTGACTGGAGTTCAGACGTGTGCTCTTCCGATCT | CAGGAAACAGCTATGAC |
| NGS #42 | [N/S/E]504 | AATGATACGGCGACCACCGAGATCTACAC | AGAGTAGA | ACACTCTTTCCCTACACGACGCTCTTCCGATCT | GGCTCT | CTTGCTTCCTGGCACGAG | N706 | CAAGCAGAAGACGGCATACGAGAT | CATGCCTA | GTGACTGGAGTTCAGACGTGTGCTCTTCCGATCT | CAGGAAACAGCTATGAC |
| NGS #43 | [N/S/E]504 | AATGATACGGCGACCACCGAGATCTACAC | AGAGTAGA | ACACTCTTTCCCTACACGACGCTCTTCCGATCT | GGCTCT | CTTGCTTCCTGGCACGAG | N707 | CAAGCAGAAGACGGCATACGAGAT | GTAGAGAG | GTGACTGGAGTTCAGACGTGTGCTCTTCCGATCT | CAGGAAACAGCTATGAC |
| NGS #44 | [N/S/E]504 | AATGATACGGCGACCACCGAGATCTACAC | AGAGTAGA | ACACTCTTTCCCTACACGACGCTCTTCCGATCT | GGCTCT | CTTGCTTCCTGGCACGAG | N708 | CAAGCAGAAGACGGCATACGAGAT | CCTCTCTG | GTGACTGGAGTTCAGACGTGTGCTCTTCCGATCT | CAGGAAACAGCTATGAC |
| NGS #45 | [N/S/E]504 | AATGATACGGCGACCACCGAGATCTACAC | AGAGTAGA | ACACTCTTTCCCTACACGACGCTCTTCCGATCT | GGCTCT | CTTGCTTCCTGGCACGAG | N709 | CAAGCAGAAGACGGCATACGAGAT | AGCGTAGC | GTGACTGGAGTTCAGACGTGTGCTCTTCCGATCT | CAGGAAACAGCTATGAC |
| NGS #46 | [N/S/E]504 | AATGATACGGCGACCACCGAGATCTACAC | AGAGTAGA | ACACTCTTTCCCTACACGACGCTCTTCCGATCT | GGCTCT | CTTGCTTCCTGGCACGAG | N710 | CAAGCAGAAGACGGCATACGAGAT | CAGCCTCG | GTGACTGGAGTTCAGACGTGTGCTCTTCCGATCT | CAGGAAACAGCTATGAC |
| NGS #47 | [N/S/E]504 | AATGATACGGCGACCACCGAGATCTACAC | AGAGTAGA | ACACTCTTTCCCTACACGACGCTCTTCCGATCT | GGCTCT | CTTGCTTCCTGGCACGAG | N711 | CAAGCAGAAGACGGCATACGAGAT | TGCCTCTT | GTGACTGGAGTTCAGACGTGTGCTCTTCCGATCT | CAGGAAACAGCTATGAC |
| NGS #48 | [N/S/E]504 | AATGATACGGCGACCACCGAGATCTACAC | AGAGTAGA | ACACTCTTTCCCTACACGACGCTCTTCCGATCT | GGCTCT | CTTGCTTCCTGGCACGAG | N712 | CAAGCAGAAGACGGCATACGAGAT | TCCTCTAC | GTGACTGGAGTTCAGACGTGTGCTCTTCCGATCT | CAGGAAACAGCTATGAC |
| NGS #49 | [N/S/E]505 | AATGATACGGCGACCACCGAGATCTACAC | GTAAGGAG | ACACTCTTTCCCTACACGACGCTCTTCCGATCT | AGGCGA | CTTGCTTCCTGGCACGAG | N701 | CAAGCAGAAGACGGCATACGAGAT | TCGCCTTA | GTGACTGGAGTTCAGACGTGTGCTCTTCCGATCT | CAGGAAACAGCTATGAC |
| NGS #50 | [N/S/E]505 | AATGATACGGCGACCACCGAGATCTACAC | GTAAGGAG | ACACTCTTTCCCTACACGACGCTCTTCCGATCT | AGGCGA | CTTGCTTCCTGGCACGAG | N702 | CAAGCAGAAGACGGCATACGAGAT | CTAGTACG | GTGACTGGAGTTCAGACGTGTGCTCTTCCGATCT | CAGGAAACAGCTATGAC |
| NGS #51 | [N/S/E]505 | AATGATACGGCGACCACCGAGATCTACAC | GTAAGGAG | ACACTCTTTCCCTACACGACGCTCTTCCGATCT | AGGCGA | CTTGCTTCCTGGCACGAG | N703 | CAAGCAGAAGACGGCATACGAGAT | TTCTGCCT | GTGACTGGAGTTCAGACGTGTGCTCTTCCGATCT | CAGGAAACAGCTATGAC |
| NGS #52 | [N/S/E]505 | AATGATACGGCGACCACCGAGATCTACAC | GTAAGGAG | ACACTCTTTCCCTACACGACGCTCTTCCGATCT | AGGCGA | CTTGCTTCCTGGCACGAG | N704 | CAAGCAGAAGACGGCATACGAGAT | GCTCAGGA | GTGACTGGAGTTCAGACGTGTGCTCTTCCGATCT | CAGGAAACAGCTATGAC |
| NGS #53 | [N/S/E]505 | AATGATACGGCGACCACCGAGATCTACAC | GTAAGGAG | ACACTCTTTCCCTACACGACGCTCTTCCGATCT | AGGCGA | CTTGCTTCCTGGCACGAG | N705 | CAAGCAGAAGACGGCATACGAGAT | AGGAGTCC | GTGACTGGAGTTCAGACGTGTGCTCTTCCGATCT | CAGGAAACAGCTATGAC |
| NGS #54 | [N/S/E]505 | AATGATACGGCGACCACCGAGATCTACAC | GTAAGGAG | ACACTCTTTCCCTACACGACGCTCTTCCGATCT | AGGCGA | CTTGCTTCCTGGCACGAG | N706 | CAAGCAGAAGACGGCATACGAGAT | CATGCCTA | GTGACTGGAGTTCAGACGTGTGCTCTTCCGATCT | CAGGAAACAGCTATGAC |
| NGS #55 | [N/S/E]505 | AATGATACGGCGACCACCGAGATCTACAC | GTAAGGAG | ACACTCTTTCCCTACACGACGCTCTTCCGATCT | AGGCGA | CTTGCTTCCTGGCACGAG | N707 | CAAGCAGAAGACGGCATACGAGAT | GTAGAGAG | GTGACTGGAGTTCAGACGTGTGCTCTTCCGATCT | CAGGAAACAGCTATGAC |
| NGS #56 | [N/S/E]505 | AATGATACGGCGACCACCGAGATCTACAC | GTAAGGAG | ACACTCTTTCCCTACACGACGCTCTTCCGATCT | AGGCGA | CTTGCTTCCTGGCACGAG | N708 | CAAGCAGAAGACGGCATACGAGAT | CCTCTCTG | GTGACTGGAGTTCAGACGTGTGCTCTTCCGATCT | CAGGAAACAGCTATGAC |
| NGS #57 | [N/S/E]505 | AATGATACGGCGACCACCGAGATCTACAC | GTAAGGAG | ACACTCTTTCCCTACACGACGCTCTTCCGATCT | AGGCGA | CTTGCTTCCTGGCACGAG | N709 | CAAGCAGAAGACGGCATACGAGAT | AGCGTAGC | GTGACTGGAGTTCAGACGTGTGCTCTTCCGATCT | CAGGAAACAGCTATGAC |
| NGS #58 | [N/S/E]505 | AATGATACGGCGACCACCGAGATCTACAC | GTAAGGAG | ACACTCTTTCCCTACACGACGCTCTTCCGATCT | AGGCGA | CTTGCTTCCTGGCACGAG | N710 | CAAGCAGAAGACGGCATACGAGAT | CAGCCTCG | GTGACTGGAGTTCAGACGTGTGCTCTTCCGATCT | CAGGAAACAGCTATGAC |
| NGS #59 | [N/S/E]505 | AATGATACGGCGACCACCGAGATCTACAC | GTAAGGAG | ACACTCTTTCCCTACACGACGCTCTTCCGATCT | AGGCGA | CTTGCTTCCTGGCACGAG | N711 | CAAGCAGAAGACGGCATACGAGAT | TGCCTCTT | GTGACTGGAGTTCAGACGTGTGCTCTTCCGATCT | CAGGAAACAGCTATGAC |
| NGS #60 | [N/S/E]505 | AATGATACGGCGACCACCGAGATCTACAC | GTAAGGAG | ACACTCTTTCCCTACACGACGCTCTTCCGATCT | AGGCGA | CTTGCTTCCTGGCACGAG | N712 | CAAGCAGAAGACGGCATACGAGAT | TCCTCTAC | GTGACTGGAGTTCAGACGTGTGCTCTTCCGATCT | CAGGAAACAGCTATGAC |
| NGS #61 | [N/S/E]506 | AATGATACGGCGACCACCGAGATCTACAC | ACTGCATA | ACACTCTTTCCCTACACGACGCTCTTCCGATCT | TAATCT | CTTGCTTCCTGGCACGAG | N701 | CAAGCAGAAGACGGCATACGAGAT | TCGCCTTA | GTGACTGGAGTTCAGACGTGTGCTCTTCCGATCT | CAGGAAACAGCTATGAC |
| NGS #62 | [N/S/E]506 | AATGATACGGCGACCACCGAGATCTACAC | ACTGCATA | ACACTCTTTCCCTACACGACGCTCTTCCGATCT | TAATCT | CTTGCTTCCTGGCACGAG | N702 | CAAGCAGAAGACGGCATACGAGAT | CTAGTACG | GTGACTGGAGTTCAGACGTGTGCTCTTCCGATCT | CAGGAAACAGCTATGAC |
| NGS #63 | [N/S/E]506 | AATGATACGGCGACCACCGAGATCTACAC | ACTGCATA | ACACTCTTTCCCTACACGACGCTCTTCCGATCT | TAATCT | CTTGCTTCCTGGCACGAG | N703 | CAAGCAGAAGACGGCATACGAGAT | TTCTGCCT | GTGACTGGAGTTCAGACGTGTGCTCTTCCGATCT | CAGGAAACAGCTATGAC |
| NGS #64 | [N/S/E]506 | AATGATACGGCGACCACCGAGATCTACAC | ACTGCATA | ACACTCTTTCCCTACACGACGCTCTTCCGATCT | TAATCT | CTTGCTTCCTGGCACGAG | N704 | CAAGCAGAAGACGGCATACGAGAT | GCTCAGGA | GTGACTGGAGTTCAGACGTGTGCTCTTCCGATCT | CAGGAAACAGCTATGAC |
| NGS #65 | [N/S/E]506 | AATGATACGGCGACCACCGAGATCTACAC | ACTGCATA | ACACTCTTTCCCTACACGACGCTCTTCCGATCT | TAATCT | CTTGCTTCCTGGCACGAG | N705 | CAAGCAGAAGACGGCATACGAGAT | AGGAGTCC | GTGACTGGAGTTCAGACGTGTGCTCTTCCGATCT | CAGGAAACAGCTATGAC |
| NGS #66 | [N/S/E]506 | AATGATACGGCGACCACCGAGATCTACAC | ACTGCATA | ACACTCTTTCCCTACACGACGCTCTTCCGATCT | TAATCT | CTTGCTTCCTGGCACGAG | N706 | CAAGCAGAAGACGGCATACGAGAT | CATGCCTA | GTGACTGGAGTTCAGACGTGTGCTCTTCCGATCT | CAGGAAACAGCTATGAC |

| | | | | | | | | | | | |
|---|---|---|---|---|---|---|---|---|---|---|---|
| **NGS #67** | **[N/S/E]506** | AATGATACGGCGACCACCGAGATCTACAC | ACTGCATA | ACACTCTTTCCCTACACGACGCTCTTCCGATCT | TAATCT | CTTGCTTCCTGGCACGAG | **N707** | CAAGCAGAAGACGGCATACGAGAT | GTAGAGAG | GTGACTGGAGTTCAGACGTGTGCTCTTCCGATCT | CAGGAAACAGCTATGAC |
| **NGS #68** | **[N/S/E]506** | AATGATACGGCGACCACCGAGATCTACAC | ACTGCATA | ACACTCTTTCCCTACACGACGCTCTTCCGATCT | TAATCT | CTTGCTTCCTGGCACGAG | **N708** | CAAGCAGAAGACGGCATACGAGAT | CCTCTCTG | GTGACTGGAGTTCAGACGTGTGCTCTTCCGATCT | CAGGAAACAGCTATGAC |
| **NGS #69** | **[N/S/E]506** | AATGATACGGCGACCACCGAGATCTACAC | ACTGCATA | ACACTCTTTCCCTACACGACGCTCTTCCGATCT | TAATCT | CTTGCTTCCTGGCACGAG | **N709** | CAAGCAGAAGACGGCATACGAGAT | AGCGTAGC | GTGACTGGAGTTCAGACGTGTGCTCTTCCGATCT | CAGGAAACAGCTATGAC |
| **NGS #70** | **[N/S/E]506** | AATGATACGGCGACCACCGAGATCTACAC | ACTGCATA | ACACTCTTTCCCTACACGACGCTCTTCCGATCT | TAATCT | CTTGCTTCCTGGCACGAG | **N710** | CAAGCAGAAGACGGCATACGAGAT | CAGCCTCG | GTGACTGGAGTTCAGACGTGTGCTCTTCCGATCT | CAGGAAACAGCTATGAC |
| **NGS #71** | **[N/S/E]506** | AATGATACGGCGACCACCGAGATCTACAC | ACTGCATA | ACACTCTTTCCCTACACGACGCTCTTCCGATCT | TAATCT | CTTGCTTCCTGGCACGAG | **N711** | CAAGCAGAAGACGGCATACGAGAT | TGCCTCTT | GTGACTGGAGTTCAGACGTGTGCTCTTCCGATCT | CAGGAAACAGCTATGAC |
| **NGS #72** | **[N/S/E]506** | AATGATACGGCGACCACCGAGATCTACAC | ACTGCATA | ACACTCTTTCCCTACACGACGCTCTTCCGATCT | TAATCT | CTTGCTTCCTGGCACGAG | **N712** | CAAGCAGAAGACGGCATACGAGAT | TCCTCTAC | GTGACTGGAGTTCAGACGTGTGCTCTTCCGATCT | CAGGAAACAGCTATGAC |
| **NGS #73** | **[N/S/E]507** | AATGATACGGCGACCACCGAGATCTACAC | AAGGAGTA | ACACTCTTTCCCTACACGACGCTCTTCCGATCT | CAGGAC | CTTGCTTCCTGGCACGAG | **N701** | CAAGCAGAAGACGGCATACGAGAT | TCGCCTTA | GTGACTGGAGTTCAGACGTGTGCTCTTCCGATCT | CAGGAAACAGCTATGAC |
| **NGS #74** | **[N/S/E]507** | AATGATACGGCGACCACCGAGATCTACAC | AAGGAGTA | ACACTCTTTCCCTACACGACGCTCTTCCGATCT | CAGGAC | CTTGCTTCCTGGCACGAG | **N702** | CAAGCAGAAGACGGCATACGAGAT | CTAGTACG | GTGACTGGAGTTCAGACGTGTGCTCTTCCGATCT | CAGGAAACAGCTATGAC |
| **NGS #75** | **[N/S/E]507** | AATGATACGGCGACCACCGAGATCTACAC | AAGGAGTA | ACACTCTTTCCCTACACGACGCTCTTCCGATCT | CAGGAC | CTTGCTTCCTGGCACGAG | **N703** | CAAGCAGAAGACGGCATACGAGAT | TTCTGCCT | GTGACTGGAGTTCAGACGTGTGCTCTTCCGATCT | CAGGAAACAGCTATGAC |
| **NGS #76** | **[N/S/E]507** | AATGATACGGCGACCACCGAGATCTACAC | AAGGAGTA | ACACTCTTTCCCTACACGACGCTCTTCCGATCT | CAGGAC | CTTGCTTCCTGGCACGAG | **N704** | CAAGCAGAAGACGGCATACGAGAT | GCTCAGGA | GTGACTGGAGTTCAGACGTGTGCTCTTCCGATCT | CAGGAAACAGCTATGAC |
| **NGS #77** | **[N/S/E]507** | AATGATACGGCGACCACCGAGATCTACAC | AAGGAGTA | ACACTCTTTCCCTACACGACGCTCTTCCGATCT | CAGGAC | CTTGCTTCCTGGCACGAG | **N705** | CAAGCAGAAGACGGCATACGAGAT | AGGAGTCC | GTGACTGGAGTTCAGACGTGTGCTCTTCCGATCT | CAGGAAACAGCTATGAC |
| **NGS #78** | **[N/S/E]507** | AATGATACGGCGACCACCGAGATCTACAC | AAGGAGTA | ACACTCTTTCCCTACACGACGCTCTTCCGATCT | CAGGAC | CTTGCTTCCTGGCACGAG | **N706** | CAAGCAGAAGACGGCATACGAGAT | CATGCCTA | GTGACTGGAGTTCAGACGTGTGCTCTTCCGATCT | CAGGAAACAGCTATGAC |
| **NGS #79** | **[N/S/E]507** | AATGATACGGCGACCACCGAGATCTACAC | AAGGAGTA | ACACTCTTTCCCTACACGACGCTCTTCCGATCT | CAGGAC | CTTGCTTCCTGGCACGAG | **N707** | CAAGCAGAAGACGGCATACGAGAT | GTAGAGAG | GTGACTGGAGTTCAGACGTGTGCTCTTCCGATCT | CAGGAAACAGCTATGAC |
| **NGS #80** | **[N/S/E]507** | AATGATACGGCGACCACCGAGATCTACAC | AAGGAGTA | ACACTCTTTCCCTACACGACGCTCTTCCGATCT | CAGGAC | CTTGCTTCCTGGCACGAG | **N708** | CAAGCAGAAGACGGCATACGAGAT | CCTCTCTG | GTGACTGGAGTTCAGACGTGTGCTCTTCCGATCT | CAGGAAACAGCTATGAC |
| **NGS #81** | **[N/S/E]507** | AATGATACGGCGACCACCGAGATCTACAC | AAGGAGTA | ACACTCTTTCCCTACACGACGCTCTTCCGATCT | CAGGAC | CTTGCTTCCTGGCACGAG | **N709** | CAAGCAGAAGACGGCATACGAGAT | AGCGTAGC | GTGACTGGAGTTCAGACGTGTGCTCTTCCGATCT | CAGGAAACAGCTATGAC |
| **NGS #82** | **[N/S/E]507** | AATGATACGGCGACCACCGAGATCTACAC | AAGGAGTA | ACACTCTTTCCCTACACGACGCTCTTCCGATCT | CAGGAC | CTTGCTTCCTGGCACGAG | **N710** | CAAGCAGAAGACGGCATACGAGAT | CAGCCTCG | GTGACTGGAGTTCAGACGTGTGCTCTTCCGATCT | CAGGAAACAGCTATGAC |
| **NGS #83** | **[N/S/E]507** | AATGATACGGCGACCACCGAGATCTACAC | AAGGAGTA | ACACTCTTTCCCTACACGACGCTCTTCCGATCT | CAGGAC | CTTGCTTCCTGGCACGAG | **N711** | CAAGCAGAAGACGGCATACGAGAT | TGCCTCTT | GTGACTGGAGTTCAGACGTGTGCTCTTCCGATCT | CAGGAAACAGCTATGAC |
| **NGS #84** | **[N/S/E]507** | AATGATACGGCGACCACCGAGATCTACAC | AAGGAGTA | ACACTCTTTCCCTACACGACGCTCTTCCGATCT | CAGGAC | CTTGCTTCCTGGCACGAG | **N712** | CAAGCAGAAGACGGCATACGAGAT | TCCTCTAC | GTGACTGGAGTTCAGACGTGTGCTCTTCCGATCT | CAGGAAACAGCTATGAC |
| **NGS #85** | **[N/S/E]508** | AATGATACGGCGACCACCGAGATCTACAC | CTAAGCCT | ACACTCTTTCCCTACACGACGCTCTTCCGATCT | GTACTG | CTTGCTTCCTGGCACGAG | **N701** | CAAGCAGAAGACGGCATACGAGAT | TCGCCTTA | GTGACTGGAGTTCAGACGTGTGCTCTTCCGATCT | CAGGAAACAGCTATGAC |
| **NGS #86** | **[N/S/E]508** | AATGATACGGCGACCACCGAGATCTACAC | CTAAGCCT | ACACTCTTTCCCTACACGACGCTCTTCCGATCT | GTACTG | CTTGCTTCCTGGCACGAG | **N702** | CAAGCAGAAGACGGCATACGAGAT | CTAGTACG | GTGACTGGAGTTCAGACGTGTGCTCTTCCGATCT | CAGGAAACAGCTATGAC |
| **NGS #87** | **[N/S/E]508** | AATGATACGGCGACCACCGAGATCTACAC | CTAAGCCT | ACACTCTTTCCCTACACGACGCTCTTCCGATCT | GTACTG | CTTGCTTCCTGGCACGAG | **N703** | CAAGCAGAAGACGGCATACGAGAT | TTCTGCCT | GTGACTGGAGTTCAGACGTGTGCTCTTCCGATCT | CAGGAAACAGCTATGAC |
| **NGS #88** | **[N/S/E]508** | AATGATACGGCGACCACCGAGATCTACAC | CTAAGCCT | ACACTCTTTCCCTACACGACGCTCTTCCGATCT | GTACTG | CTTGCTTCCTGGCACGAG | **N704** | CAAGCAGAAGACGGCATACGAGAT | GCTCAGGA | GTGACTGGAGTTCAGACGTGTGCTCTTCCGATCT | CAGGAAACAGCTATGAC |
| **NGS #89** | **[N/S/E]508** | AATGATACGGCGACCACCGAGATCTACAC | CTAAGCCT | ACACTCTTTCCCTACACGACGCTCTTCCGATCT | GTACTG | CTTGCTTCCTGGCACGAG | **N705** | CAAGCAGAAGACGGCATACGAGAT | AGGAGTCC | GTGACTGGAGTTCAGACGTGTGCTCTTCCGATCT | CAGGAAACAGCTATGAC |
| **NGS #90** | **[N/S/E]508** | AATGATACGGCGACCACCGAGATCTACAC | CTAAGCCT | ACACTCTTTCCCTACACGACGCTCTTCCGATCT | GTACTG | CTTGCTTCCTGGCACGAG | **N706** | CAAGCAGAAGACGGCATACGAGAT | CATGCCTA | GTGACTGGAGTTCAGACGTGTGCTCTTCCGATCT | CAGGAAACAGCTATGAC |
| **NGS #91** | **[N/S/E]508** | AATGATACGGCGACCACCGAGATCTACAC | CTAAGCCT | ACACTCTTTCCCTACACGACGCTCTTCCGATCT | GTACTG | CTTGCTTCCTGGCACGAG | **N707** | CAAGCAGAAGACGGCATACGAGAT | GTAGAGAG | GTGACTGGAGTTCAGACGTGTGCTCTTCCGATCT | CAGGAAACAGCTATGAC |
| **NGS #92** | **[N/S/E]508** | AATGATACGGCGACCACCGAGATCTACAC | CTAAGCCT | ACACTCTTTCCCTACACGACGCTCTTCCGATCT | GTACTG | CTTGCTTCCTGGCACGAG | **N708** | CAAGCAGAAGACGGCATACGAGAT | CCTCTCTG | GTGACTGGAGTTCAGACGTGTGCTCTTCCGATCT | CAGGAAACAGCTATGAC |
| **NGS #93** | **[N/S/E]508** | AATGATACGGCGACCACCGAGATCTACAC | CTAAGCCT | ACACTCTTTCCCTACACGACGCTCTTCCGATCT | GTACTG | CTTGCTTCCTGGCACGAG | **N709** | CAAGCAGAAGACGGCATACGAGAT | AGCGTAGC | GTGACTGGAGTTCAGACGTGTGCTCTTCCGATCT | CAGGAAACAGCTATGAC |
| **NGS #94** | **[N/S/E]508** | AATGATACGGCGACCACCGAGATCTACAC | CTAAGCCT | ACACTCTTTCCCTACACGACGCTCTTCCGATCT | GTACTG | CTTGCTTCCTGGCACGAG | **N710** | CAAGCAGAAGACGGCATACGAGAT | CAGCCTCG | GTGACTGGAGTTCAGACGTGTGCTCTTCCGATCT | CAGGAAACAGCTATGAC |
| **NGS #95** | **[N/S/E]508** | AATGATACGGCGACCACCGAGATCTACAC | CTAAGCCT | ACACTCTTTCCCTACACGACGCTCTTCCGATCT | GTACTG | CTTGCTTCCTGGCACGAG | **N711** | CAAGCAGAAGACGGCATACGAGAT | TGCCTCTT | GTGACTGGAGTTCAGACGTGTGCTCTTCCGATCT | CAGGAAACAGCTATGAC |
| **NGS #96** | **[N/S/E]508** | AATGATACGGCGACCACCGAGATCTACAC | CTAAGCCT | ACACTCTTTCCCTACACGACGCTCTTCCGATCT | GTACTG | CTTGCTTCCTGGCACGAG | **N712** | CAAGCAGAAGACGGCATACGAGAT | TCCTCTAC | GTGACTGGAGTTCAGACGTGTGCTCTTCCGATCT | CAGGAAACAGCTATGAC |

Illumina adapters N701-N712 and [N/S/E]501-[N/S/E]508 and [N/S/E]517) were used, which lead to 96 different combinations of primer pairs. Through the addition of these adapter primers additional DNA motifs are introduced, such as index and variable sequences, sequencing primer binding sites and regions complementary to the flow cell Oligos which are critical for cluster generation. The combination of index and variale sequences in forward and reverse primers are unique to each set of primer pair and serve as identifiers for each study subject. As they  are specific to a given sample library they enable multiple sequences to be sequenced together and are used for de-multiplexing during data analysis to assign individual sequence reads to the correct sample during final data analysis.

**Appendix 5: Description of the study cohort**

| | BPD | | HC | | T Test | |
|---|---|---|---|---|---|---|
| | *M* | *SD* | *M* | *SD* | *t* | *p* |
| Age | 26.3 | 5.7 | 24.5 | 4.4 | -1.72 | 0.089 |
| IQ | 108.0 | 15.0 | 111.2 | 14.6 | 1.03 | 0.307 |
| Childhood Trauma Questionnaire | | | | | | |
| total score | 66.0 | 19.9 | 33.1 | 9.7 | -9.95 | < 0.001 |
| emotional abuse | 18.0 | 5.7 | 7.4 | 3.3 | -10.77 | < 0.001 |
| physical abuse | 9.6 | 4.4 | 5.7 | 1.6 | -5.54 | < 0.001 |
| sexual abuse | 10.5 | 6.0 | 6.0 | 1.5 | -5.53 | < 0.001 |
| emotional neglect | 17.6 | 5.9 | 8.4 | 3.3 | -9.03 | < 0.001 |
| physical neglect | 10.4 | 4.5 | 6.3 | 1.9 | -5.66 | < 0.001 |
| Interpersonal Reactivity Index | | | | | | |
| perspective taking | 13.0 | 6.9 | 18.8 | 4.0 | 4.99 | < 0.001 |
| fantasy | 17.0 | 6.9 | 18.8 | 5.0 | 1.38 | 0.172 |
| empathic concern | 18.9 | 6.3 | 19.8 | 4.1 | 0.81 | 0.420 |
| personal distress | 21.8 | 4.6 | 12.5 | 4.3 | -9.91 | < 0.001 |

Patients with borderline personality disorder (BPD) compared to healthy controls (HC)

A)

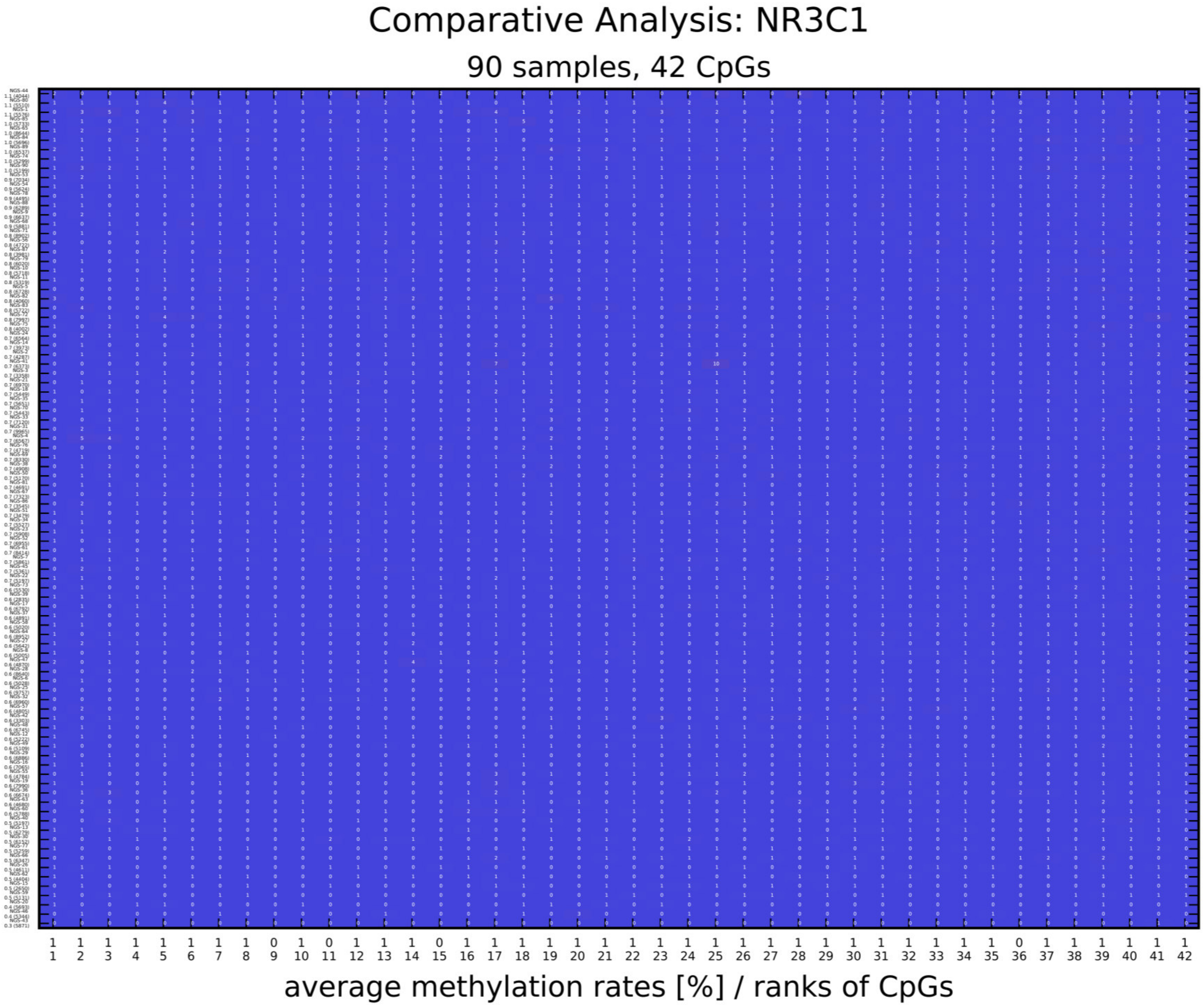
Comparative Analysis: NR3C1
90 samples, 42 CpGs
average methylation rates [%] / ranks of CpGs

B)

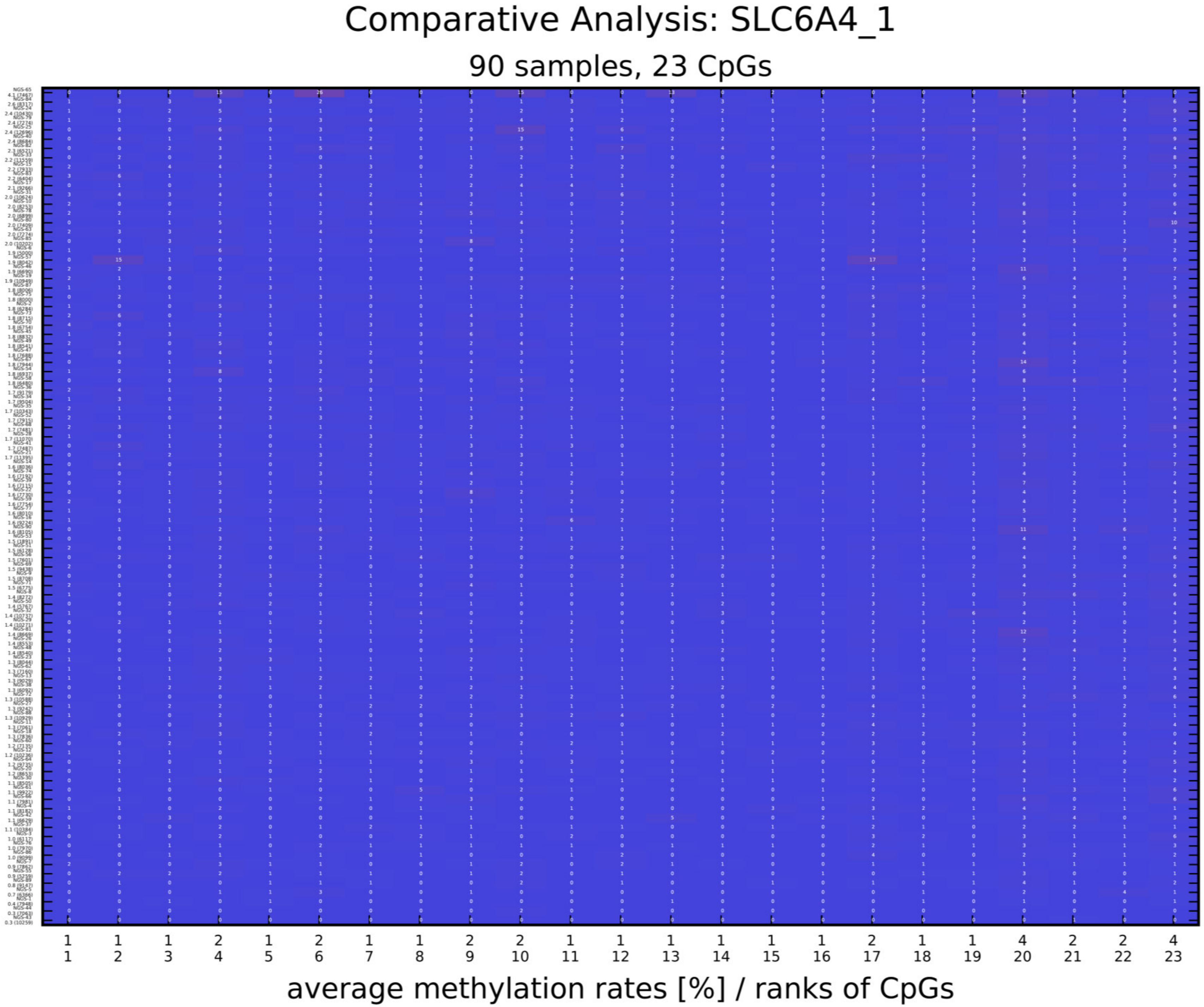
Comparative Analysis: SLC6A4_1
90 samples, 23 CpGs
average methylation rates [%] / ranks of CpGs
1 1 1 2 1 2 1 1 2 2 1 1 1 1 1 1 2 1 1 4 2 2 4
1 2 3 4 5 6 7 8 9 10 11 12 13 14 15 16 17 18 19 20 21 22 23

c)

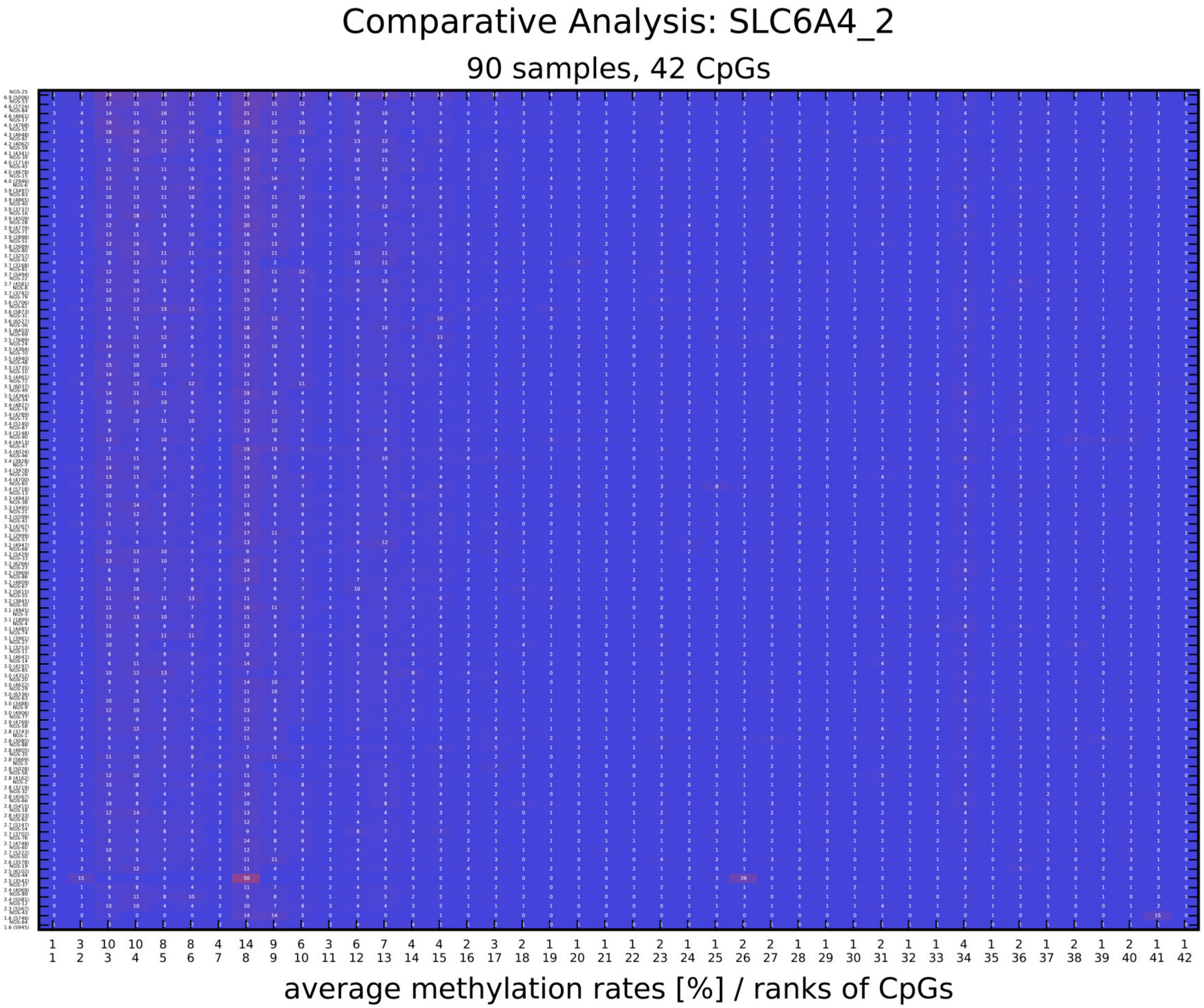

Comparative Analysis: SLC6A4_2
90 samples, 42 CpGs
1 3 10 10 8 8 4 14 9 6 3 6 7 4 4 2 3 2 1 1 1 1 2 1 1 2 2 1 1 1 2 1 1 4 1 2 1 2 1 2 1 1
1 2 3 4 5 6 7 8 9 10 11 12 13 14 15 16 17 18 19 20 21 22 23 24 25 26 27 28 29 30 31 32 33 34 35 36 37 38 39 40 41 42
average methylation rates [%] / ranks of CpGs

D)

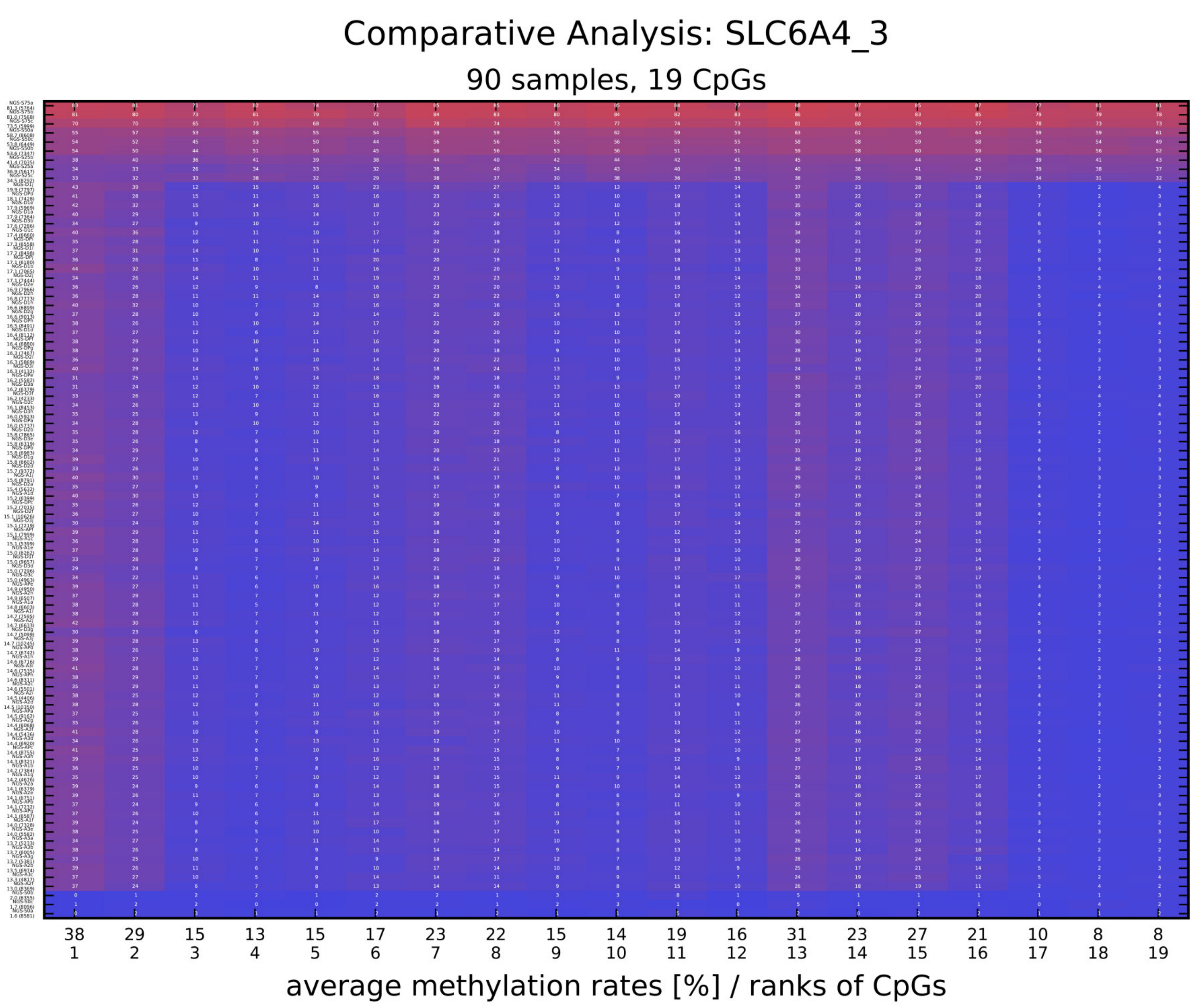

Comparative Analysis: SLC6A4_3
90 samples, 19 CpGs
38 29 15 13 15 17 23 22 15 14 19 16 31 23 27 21 10 8 8
1 2 3 4 5 6 7 8 9 10 11 12 13 14 15 16 17 18 19
average methylation rates [%] / ranks of CpGs

E)

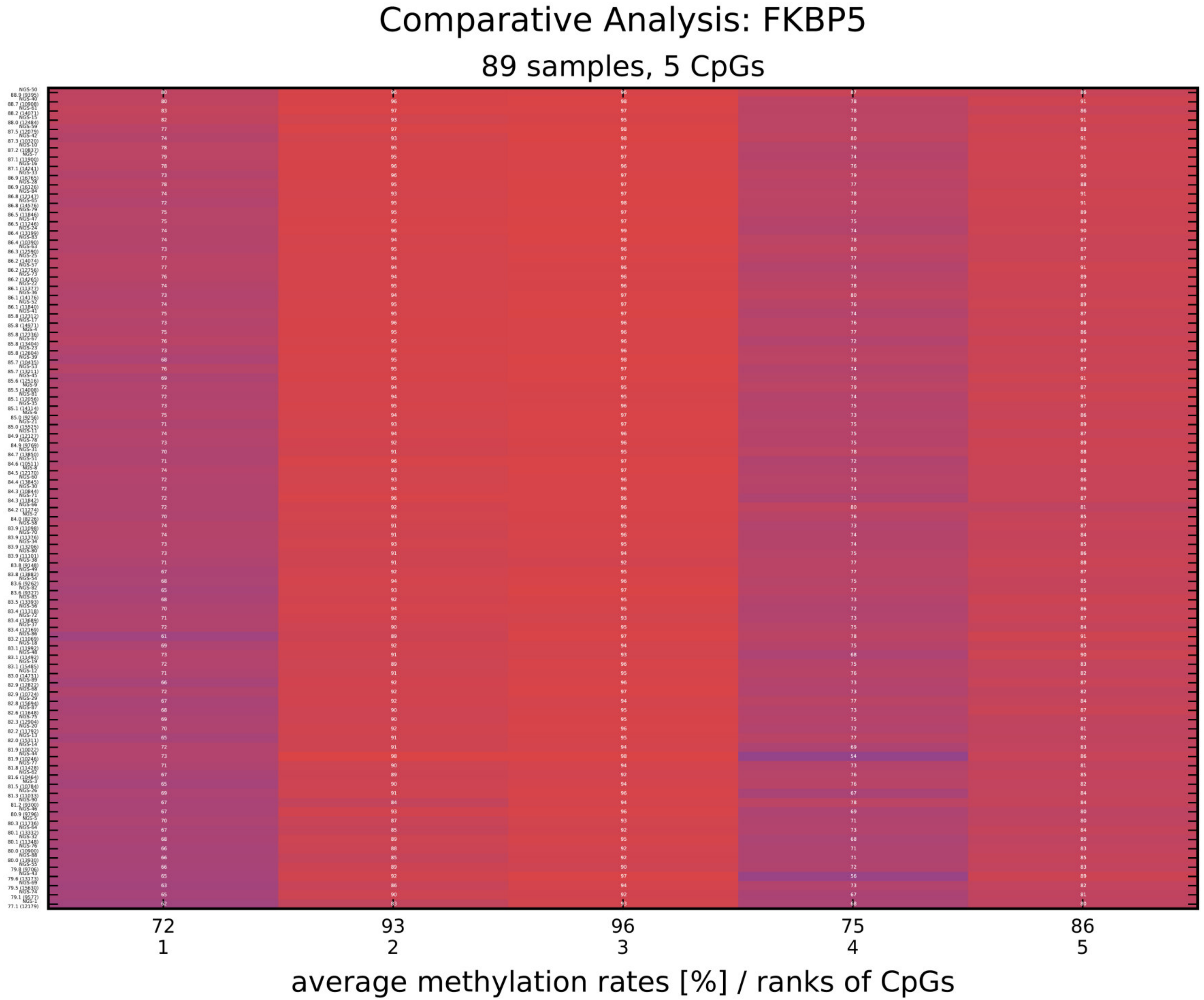

Comparative Analysis: FKBP5
89 samples, 5 CpGs
72
1
93
2
96
3
75
4
86
5
average methylation rates [%] / ranks of CpGs

F)

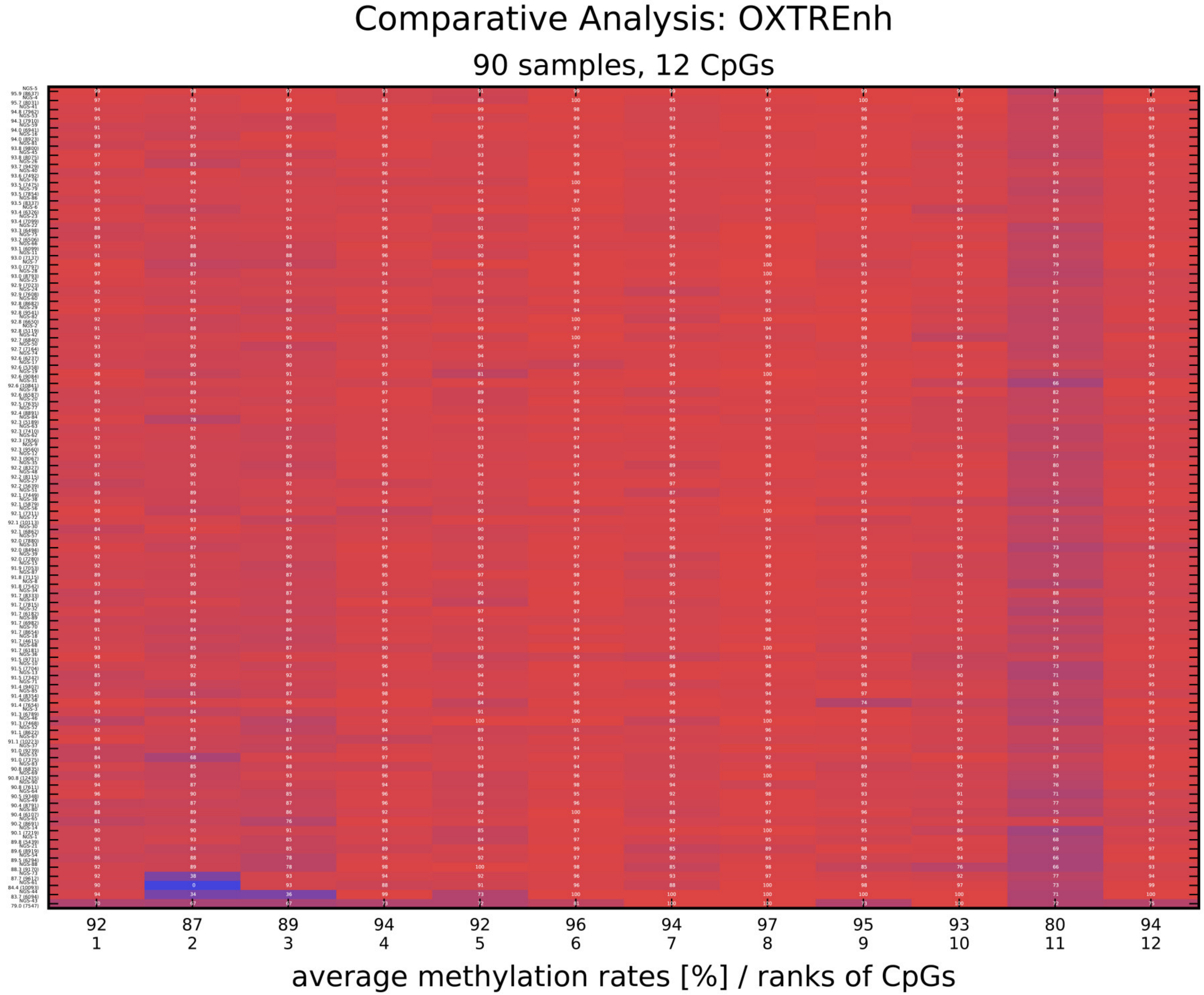
Comparative Analysis: OXTREnh
90 samples, 12 CpGs
92 87 89 94 92 96 94 97 95 93 80 94
1 2 3 4 5 6 7 8 9 10 11 12
average methylation rates [%] / ranks of CpGs

**Appendix 6: Graphical output generated by Amplikyzer2 software**

Graphical output was used to check for potential technical issues and missing sequence information, which are indicated by bars of grey color; blue color represents unmethylated CpGs and red color indicates methylated CpGs. Each line represents one sample, each column one CpG site. Average methylation rates (in %) are presented for each amplicon, including all CpGs of all subjects.

**Appendix 7: Reliability of the assays**

| | Bisulfit 1 | | Bisulfit 2 | | Bisulfit 3 | | Pool | | mean | | mean |
|---|---|---|---|---|---|---|---|---|---|---|---|
| ***NR3C1*** | **male** | **female** | **male** | **female** | **male** | **female** | **male** | **female** | **male** | **female** | **all** |
| mean SD | 0.98 | 0.53 | 0.89 | 0.4 | 1.0 | 0.28 | 0.67 | 0.62 | 0.88 | 0.46 | 0.67 |
| max SD | (2.26) | (0.84) | (2.15) | (2.7) | (2.7) | (1.63) | (1.56) | (1.25) | (2.7) | (2.7) | |
| ***SLC6A4_1*** | | | | | | | | | | | |
| mean SD | 0.92 | 1.01 | 0.99 | 2.26 | 1.22 | 1.84 | 1.10 | 1.64 | 1.06 | 1.69 | 1.37 |
| max SD | (1.95) | (2.16) | (1.54) | (3.70) | (1.93) | (3.09) | (1.95) | (2.56) | (1.95) | (3.7) | |
| ***SLC6A4_2*** | | | | | | | | | | | |
| mean SD | 1.07 | 1.46 | 1.23 | 2.62 | 1.63 | 2.23 | 1.37 | 1.48 | 1.31 | 1.48 | 1.39 |
| max SD | (2.28) | (3.22) | (2.60) | (6.10) | (2.92) | (5.85) | (3.19) | (3.11) | (3.19) | 6.10) | |
| ***SLC6A4_3*** | | | | | | | | | | | |
| mean SD | 2.11 | 1.24 | 1.58 | 1.37 | 1.89 | 1.62 | 1.56 | 1.24 | 1.78 | 1.24 | 1.51 |
| max SD | (3.22) | (2.06) | (2.33) | (2.16) | (2.83) | (2.88) | (3.16) | (2.05) | (3.22) | (2.05) | |
| ***FKBP5*** | | | | | | | | | | | |
| mean SD | 1.53 | 0.94 | 1.55 | 1.51 | 1.63 | 1.04 | 1.26 | 0.90 | 1.49 | 1.09 | 1.29 |
| max SD | (2.77) | (1.59) | (2.27) | (1.89) | (2.46) | (1.81) | (1.77) | (1.14) | (2.77) | (1.89) | |
| ***OXTR_Enh*** | | | | | | | | | | | |
| mean SD | 2.00 | 2.19 | 1.50 | 2.96 | 2.35 | n.a. | 1.96 | 3.28 | 1.91 | 2.57 | 2.24 |
| max SD | (3.48) | (3.32) | (2.55) | (3.76) | (3.23) | n.a. | (2.82) | (5.55) | (3.23) | (3.76) | |

DNA from a male and a female donor was each bisulfite-treated in three independent trials. From each trial and from a pool out of the three 10 independently generated PCR-replicates were sequenced. Mean standard deviation in percent and maximum (SD) for all CpGs per amplicon are displayed.

### FKBP5

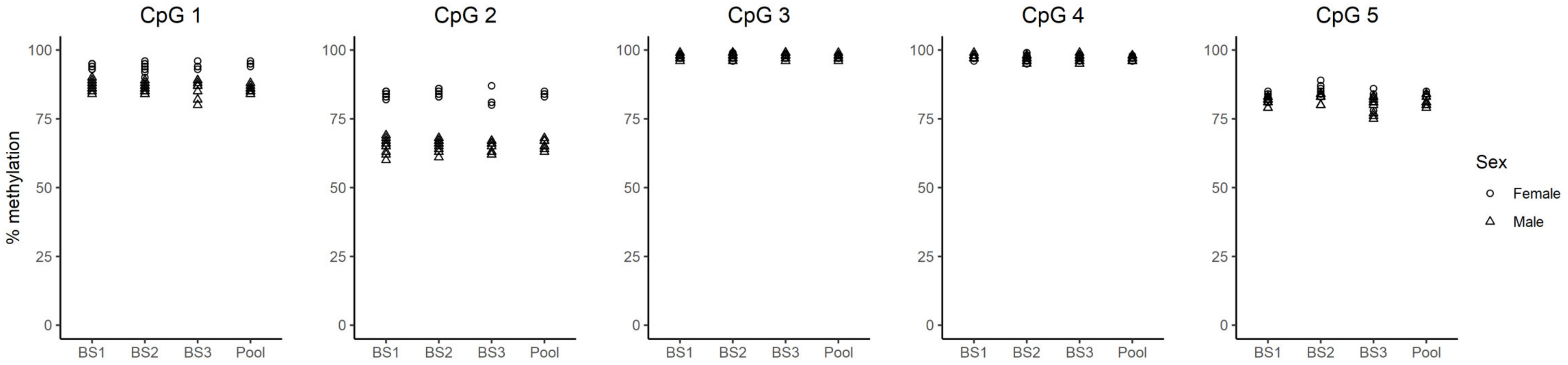


**Appendix 8: Inter-individual differences in DNA methylation**
DNA methylation analysis of DNA from a female and a male donor revealed specific differences in DNA methylation at one CpG site (CpG2). Larger deviations as also detectable at CpG sites 1 and 5 might be attributed to inter-individual differences and potentially are not based on the accuracy of the quantification method.
